# Contralesional hippocampal spreading depolarization promotes functional recovery after stroke

**DOI:** 10.1101/2023.08.31.555814

**Authors:** Andrew K.J. Boyce, Yannick Fouad, Renaud C. Gom, Donovan M. Ashby, Cristina Martins-Silva, Leonardo Molina, Tamas Füzesi, Carina Ens, Wilten Nicola, Alexander McGirr, G. Campbell Teskey, Roger J. Thompson

## Abstract

Ischemic stroke, brain tissue infarction following obstructed cerebral blood flow, leads to long-term neurological deficits and death. While neocortex is a commonly affected region with established preclinical models, less is known about deeper brain strokes, despite having unique neurological outcomes. We induced focal ischemic stroke while simultaneously monitors neuronal activity in awake behaving Thy1-GCaMP6f mice by delivering and collecting light through bilateral fiberoptic implants. Unilateral hippocampal stroke resulted in atypical mouse behavior coincident with ipsilesional terminal spreading depolarization (sustained increase in GCaMP6f fluorescence). Ischemia induced seizures that propagated to the contralesional hippocampus triggering a transient spreading depolarization (SD), predominantly in females. Hippocampal stroke impaired contextual fear conditioning acquired pre-stroke. Yet, 7 days post-stroke, contextual fear conditioning was only improved in mice with evidence of contralesional SD. Here, recovery of hippocampal function was lost by blunting peri-stroke SD with NMDAR antagonism, indicating that contralesional SD improves recovery following hippocampal stroke.

## Introduction

Ischemic stroke, the death of brain tissue triggered by obstructed cerebral blood flow, is second among global causes of death, leaving many survivors permanently disabled^1^. Ischemia in middle cerebral artery (MCA)-fed territory accounts for approximately half of focal human strokes^2^ and has been modelled through several approaches in animals^3^; however, many strokes occur in deeper brain structures. Stroke is less well understood in subcortical areas and has been understudied in rodent models due in part to technical limitations preventing optical and surgical access for stroke induction, particularly in awake animals. Of single large vessel strokes excluding those occurring in MCA-fed territory, approximately one-third occur in posterior cerebral artery (PCA)-fed territory, the vasculature that supplies the hippocampus^2,4^, a structure that mediates episodic and contextual memory and spatial navigation in both rodents^5^ and humans^6^.

Stroke symptoms can be focal, linked to deficits in the infarcted tissue, or non-focal, impacted by perturbations to remote tissue^1^. Patients most often present with a combination of both focal and non-focal symptoms^7,8^, and non-focal presenting symptoms may be more frequent in females (^8,9^ yet, see ^10^). Although hippocampal stroke is classified as rare, patients present with unique symptomology, including permanent or transient amnestic syndrome that can last between weeks and months^11–13^. These symptoms may be accompanied by disorientation, spatial memory deficits, and alexia without agraphia^13^. Hippocampal stroke is typically a unilateral injury^11^, resulting in complex evolution and reorganization of hippocampal networks. The mechanisms whereby the contralesional hippocampal function is impaired or preserved by this injury are poorly understood. Consequently, hippocampal stroke requires directed investigation to allow for future therapies to treat its unique neurological outcomes.

The peri-infarct period is complex with dynamic interaction of pathological and adaptive effectors^1^. Here, the dysregulated metabolic environment of focal ischemia is ripe for the initiation of additive sequelae that can radiate from injured tissue to impact network partners^14^. These include seizure, synchronous high frequency neuronal firing that can rapidly propagate across the brain via synaptic transmission (∼ 1-1000 mm/s)^15–17^, and spreading depolarizations (SD), first described by Leão^18,19^, waves of near complete neuronal and glial cell depolarization that slowly propagate through contiguous grey matter (∼ 2-5 mm/min)^20^. In severely ischemic tissue, SDs arise within ∼ 5 mins due to a supply-demand mismatch, where increased oxygen utilization is not matched by adequate perfusion^19,21–25^; this mismatch is exacerbated by ischemic sequelae including seizure. Inadequate perfusion and oxidative substrates cause ATP to drop below a critical threshold for maintaining Na^+^/K^+^-ATPase-dependent ionic gradients, arresting spontaneous activity and triggering SD, which become terminal without timely reperfusion ^26–30^, marked by the appearance of the irreversible negative ultraslow potential ^25^. Where tissue is adequately or partially perfused, SDs can drive a transient and reversible loss of spontaneous and evoked neuronal activity with a high metabolic cost to reestablish the resting membrane potential^20^. Successive and clustered SDs provide a particularly intense challenge to already metabolically challenged tissue^31^, thus penumbral tissue may succumb in the hours and days following the onset of ischemia. This results in lesion expansion, due in part to excitotoxicity and spreading oligemia and ischemia, induced by impaired neurovascular coupling and constriction of the penumbral microvasculature^31–33^. In the event of adequate perfusion or timely reperfusion following ischemia, SDs can be recovered by subsequent hyperemia that provides the energy substrate to re-establish resting membrane potential^31,32^ and allows neurons to survive ^25,26,34^. These are not distinct phenomena, but exist in a space and time continuum outwards from the ischemic core into healthy tissue^20,35^. Taken together, while it is generally accepted that SDs in ischemic core and hypoperfused (penumbral) tissue worsen stroke outcomes (^23,25,32,36^ yet, see ^37^), SDs propagating through nearby adequately-perfused tissue may not be detrimental^38–45^.

In this study, we induced unilateral ischemia in the hippocampus while simultaneously recording neuronal GCaMP6f fluorescence (proxy for intracellular Ca^2+^) in awake and freely behaving (non-head-fixed) mice. We induced unilateral hippocampal stroke via photothrombosis, a method pioneered by Rosenblum & El-Sabban^46^ and Watson *et al.*^47^, through a fiberoptic cannula implanted dorsal to the CA1 in transgenic mice expressing the Ca^2+^ biosensor GCaMP6f in pyramidal neurons (Thy1-GCaMP6f) to understand the symptomology of hippocampal ischemia and SD (Fig. 1a). While ischemia-induced SD is typically limited to the ipsilesional hemisphere during a superficial cortical ischemia^31,48^, we show that unilateral photothrombosis triggered localized terminal SD in the ipsilesional hippocampus (sustained increase in GCaMP6f). and a coincident increase in ambulation and rearing around an open field arena. Given the acute behavioural impact of the unilateral ischemia, we were interested in the impact on the well-perfused contralesional hippocampus. Here, within 10 min of terminal SD in the ischemic hippocampus, a massive transient wave of neuronal depolarization consistent with an SD also appeared in the contralesional hippocampus. Seizures are often associated with SD in both rodents^49–53^ and humans^54^. We observed contralesional electrographic seizure directly preceding SD when LFP was simultaneously recorded with neuronal Ca^2+^. Contralesional SDs were more frequent in female mice and were, surprisingly, associated with improved post-stroke contextual fear conditioning. Antagonism of N-methyl-D-aspartate receptors (NMDAR) immediately prior to unilateral photothrombosis reduced contralesional SD amplitude, worsened seizures, and abolished the beneficial outcomes on memory associated with contralesional SD. Taken together, we show that unilateral hippocampal ischemia triggers contralesional seizure followed by a transient hippocampal SD, playing a surprising protective role in contextual fear conditioning during stroke recovery, predominantly in females.

**Figure 1.**
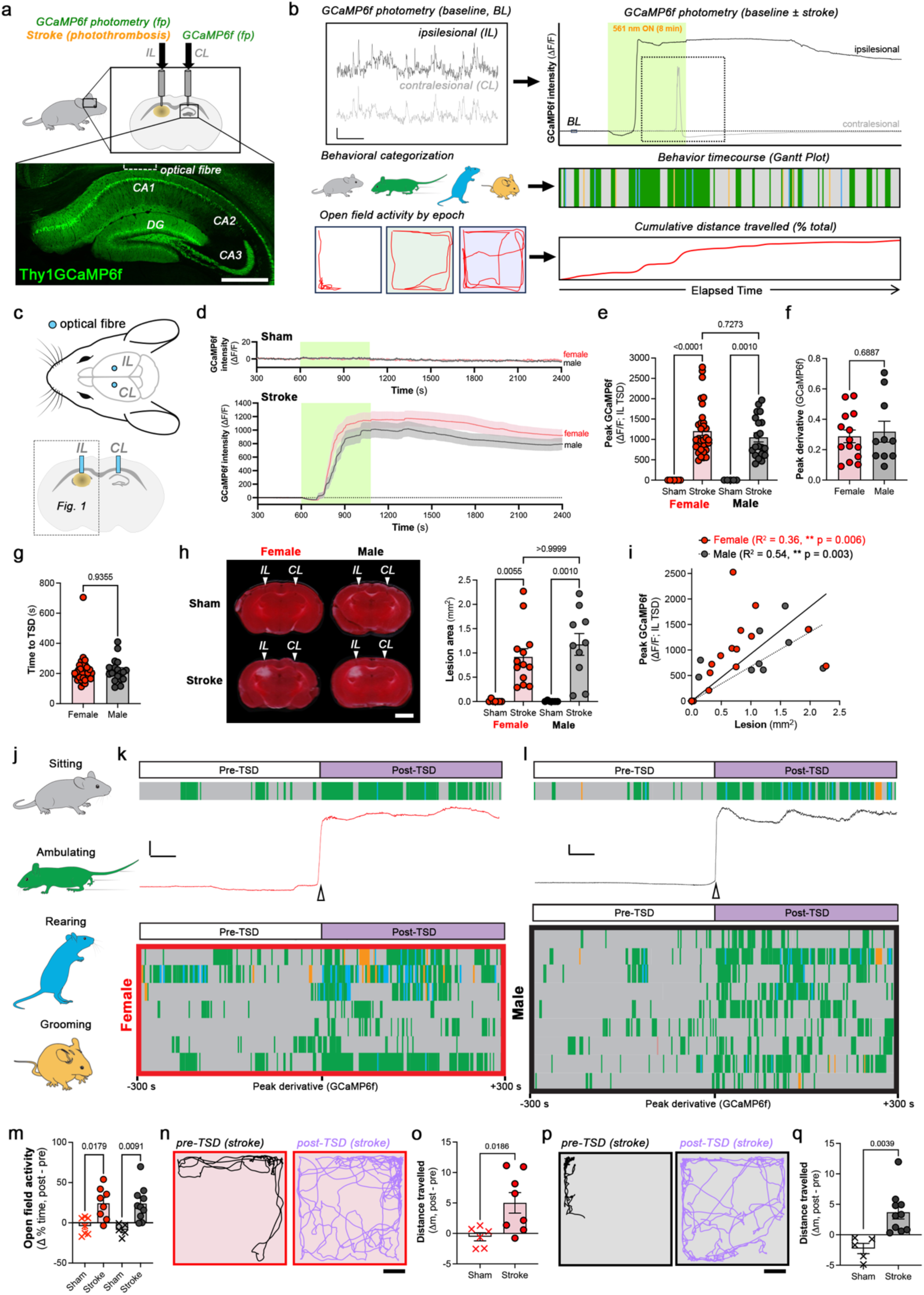
Hippocampal ischemia-evoked terminal SD correlates with lesion size and increased activity. **a** Bilateral fiberoptic cannulas for unilateral hippocampal photothrombotic stroke and bilateral photometry in awake behaving Thy1-GCaMP6f mice. **b** Basal GCaMP6f photometry (scale bar: ΔF/F = 2 arbitrary fluorescence units (arb. units), t = 5 s) ischemia-evoked GCaMP6f responses in bilateral hippocampi, paired to behavior and locomotion analyses in an open field (0.5 m^2^). Photothrombotic illumination (561 nm, 6 mW, 8 min, green box) via fiber dorsal to right CA1. **c** IL (Fig. 1) and CL (Fig. 2) effects of unilateral photothrombosis. **d** IL GCaMP6f traces (ΔF/F, mean ± s.e.m.) for female (red) and male (black) sham mice (vehicle, i.p.). IL terminal SD (TSD, ΔF/F, mean ± s.e.m.) induced by unilateral hippocampal photothrombosis in female (red) and male (black) stroke mice (1 % RB, i.p.). **e** No sex differences between IL TSD amplitude (peak ΔF/F,), **f** peak derivative of TSD, or **g** time to TSD. **h** Unilateral photothrombosis generates a focal hippocampal lesion (TTC-stained coronal sections from female (red) and male (black) sham/stroke, 48 h post-illumination; arrow highlights fiber location). Scale bars, 2 mm. No sex differences in lesion size (mm^2^). **i** IL TSD amplitude (ΔF/F) correlates with lesion size. **j** Behavior state (sitting – grey, ambulating – green, rearing – blue, and grooming – orange) during peri-TSD period (TSD ± 5 min). Behavior state (Gantt plot) aligned to representative **k** female (red) or **l** male TSDs (peak derivative, black) show coincident increase in activity at TSD onset. Scale bar: 100 arb. units, 50 s. Gantt behavioral plots for all analyzed female (red) or male (black) traces centered on the IL TSD. One mouse per row. **m** Time active (% of epoch ambulating/rearing) increases after ipsilesional TSD in both females (red, n = 7) and males (black, n = 9). In shams, behavioral traces were aligned to the average TSD onset time in stroke group (800 s; *see Extended Data* Fig 5). **n** Locomotion trace for a female before (*pre-TSD)* and after *(post-TSD)* TSD. **o** Following IL TSD, females travel further relative to shams. **p** Locomotion traces for a male pre-TSD and post-TSD. **q** Following IL TSD, males travel further relative to shams. Additional statistical information is available in Supplementary Information. Source data are provided in the Source Data file.

## Results

### Hippocampal ischemia-evoked terminal SD correlates with lesion size and increased activity

Elevated neuronal Ca^2+^ influx is a critical component of SD and the SD-initiated excitotoxic cell death in the acute phase of an ischemic stroke^55^. Here, massive influx follows the onset of the DC shift in SD^56–58^. We developed a focal stroke model in freely behaving mice for simultaneously recording both ipsilesional and contralesional neuronal Ca^2+^ dynamics in bilateral hippocampi and analyzing mouse behavior in the acute phase of unilateral hippocampal stroke using a combination of fiber photometry and photothrombosis (Fig. 1a). Thy1-GCaMP6f transgenic mice, expressing the genetically encoded Ca^2+^ indicator in pyramidal neurons, were used in our study. Here, an optical fiber (400 µm diameter) was implanted dorsal to both hippocampi (∼ 100 μm above CA1, Fig. 1a; > 10 days before sham illumination of photothrombosis) to allow photothrombosis and GCaMP6f photometry (Fig 1). After recovery and handling, mice were injected (intra-peritoneal, i.p.) with Rose Bengal (1% RB, 0.1 mg/kg) and placed into an open field where CA1 pyramidal neuron GCaMP6f fluorescence (via fiber photometry) and behavior (via overhead camera) were simultaneously recorded (Fig. 1b). To control for the fiberoptic implant, sham animals were used as a control, where fiberoptic cannulas were implanted and identical photo-illumination was performed; however, in lieu of Rose Bengal, vehicle (PBS) was injected i.p. prior to initiating the experiment. In both sham and stroke mice, a baseline was recorded (10 min) to allow unimpeded collection of behavioral and photometry data for within-mouse analyses from which to quantify ΔF/F and changes in behavior. Following baseline, laser illumination (561 nm: 8 min, 6 mW) through one optical fiber initiated the focal ischemic cascade through unilateral photo-oxidation of RB (photothrombosis, Fig. 1c). Ipsilesional photothrombosis and photometry occur through the same fiberoptic cannula, thus changes in GCaMP6f intensity in response to photothrombotic illumination represent changes in neuronal Ca^2+^ in the ischemic core. In the ipsilesional hippocampus, unilateral photothrombosis elicited a large and sustained increase in GCaMP6f intensity (up to ∼ 500-fold of basal fluctuations), which was absent in sham mice (vehicle; Fig. 1de). The onset of this sustained increase in intracellular Ca^2+^ was defined as the time when GCaMP6f intensity reached its maximum rate of change (peak derivative, Fig. 1de). Neurons undergoing sustained SD in severely ischemic tissue will die within 20 - 30 min^36^, in the absence of timely reperfusion. Given the persistent sustained increase in neuronal Ca^2+^ recorded through the ipsilesional fiber dorsal to the emerging core (> 30 min post-photothrombosis), we deemed this change in GCaMP6f intensity the terminal SD (TSD). No sex differences were observed in the amplitude of the ipsilesional TSD (Peak GCaMP6f_IL_, Fig. 1de). Further, the peak derivative of GCaMP6f (Fig. 1f) and the time between the initiation of photothrombosis and TSD (time to TSD_IL_, Fig. 1g) were also indistinguishable between sexes. Ipsilesional TSD amplitude scaled with illumination time (1.5 – 8 min; 6 mW) and laser power (4 – 6 mW; 1.5 min; Extended Data Fig. 1).

Lesions were assessed using triphenyl tetrazolium chloride (TTC), a live-dead stain relying on functional mitochondrial succinate dehydrogenase to convert a colorless substrate into a red product in metabolically active tissue (colorless in infarcted tissue, 48 h post stroke), and immunohistochemistry (7 days post stroke). Focal hippocampal stroke lesion size (TTC-negative) was indistinguishable between sexes (females: 0.92 ± 0.16 mm^2^, 1.18 ± 0.22 mm^2^; Fig. 1h). Lesions were smaller with decreased laser power and/or illumination duration (Extended Data Fig. 1). The infarct below the fiberoptic tip at 48 h post-photothrombosis (Fig. 1h) provided further support for deeming the sustained elevation of ipsilesional Ca^2+^ during our acute photometry recordings a bona fide terminal SD (TSD, Fig. 1d). We identified a correlation between TSD amplitude (ΔF/F) and lesion size in both sexes (Fig. 1i). Here, we suggest that ipsilesional TSD (GCaMP6f) could act as a complimentary non-histological approach for evaluating the impact of focal ischemia with the benefit of allowing the mouse to remain alive for long term behavioral evaluation and within-animal comparison. Of note, there was some damage to the overlying neocortex and minor glial reactivity around the implanted fiber in sham mice, which could be considered a caveat of this model; however, these effects are similar in stroke mice, hence the importance of sham surgery and illumination.

To further validate the impact of sham illumination and stroke, as well as fiberoptic implant dorsal to the hippocampus, on neuronal loss and glial reactivity, unilateral implants were made and either sham or stroke (561 nm ± 1% RB: 6 mW illumination, 8 min) was triggered. Comparisons were made between tissue in the illuminated and naïve contralateral hemisphere. Thy1-GCaMP6f mice have stable and sparse expression of GCaMP6f in layers 2/3 and 5 of the neocortex and dense expression in CA3, CA1, and the dentate gyrus of the hippocampus^59^. One week after sham or stroke illumination, coronal slices were prepared and immunolabeled for indicators of astrocyte and microglial reactivity (GFAP and Iba1, respectively) from paraformaldehyde-fixed brains extracted one week after illumination. Thus, the hippocampus was traced and the area that was GCaMP6f-positive within the hippocampus was quantified. Stroke reduced GCaMP6f in the ipsilesional hippocampus indicating neuronal loss (Extended Data Fig. 2), relative to the contralesional hippocampus, a decrease that was absent in shams and indistinguishable between sexes. GFAP and Iba1 were enriched in the ipsilesional hemisphere, surrounding the stroke core, and no sex differences were observed (Extended Data Fig. 2). Following unilateral hippocampal stroke, there was also a moderate increase in GFAP-reactivity in the contralesional hemisphere; however, this was not significantly different from the ipsilesional hemisphere nor the sham condition.

### Hippocampal stroke in awake behaving mice avoids anesthetic effects on ischemic TSD

In many existing stroke models, ischemia is initiated in the presence of anesthetic^3^. Inhaled anesthesia (i.e. isoflurane) is neuroprotective against ischemic stroke in the superficial neocortex^60–62^, complicating the interpretation of results and masking effects of potential neuroprotectants. Inhaled anesthesia also inhibits SD^61,62^ in part through competitive inhibition of NMDA receptors^63,64^. The effect of anesthesia on ischemic stroke in deeper structures was unknown. Using a single unilateral fiberoptic implant dorsal to the hippocampus, we assessed ischemia-evoked TSD in awake versus anesthetized mice. Here, following vehicle or 1% RB injection (i.p.), mice were either placed under isoflurane anesthetic (5% induction, 1% maintenance) or into the awake behaving protocol, where they were allowed to freely behave in an open field. As above, a 10-min baseline was recorded prior to photothrombosis (561 nm: 6 mW, 8 min). Here, we observed differences in basal neuronal GCaMP6f transients. Basal neuronal GCaMP6f events were identified as a transient increase in signal > 4 standard deviations above the mean separated by > 0.5 s. Awake mice had more frequent GCaMP6f events of similar amplitude relative to anesthetized mice (ΔF/F ≈ 5 a.u., Extended Fig. 3ab).

As in other stroke models^60–62^, isoflurane anesthesia was protective against focal ischemic stroke in the hippocampus (Extended Data Fig. 3). Anesthetized mice had smaller and slower TSD relative to awake mice under identical photothrombotic illumination (Extended Data Fig 3acd). The onset of ischemia-induced TSD was also delayed under anesthesia (Extended Data Fig. 3e). At 48 h after photothrombosis, stroke lesions were smaller in anesthetized relative to awake mice (Extended Data Fig. 3f). Further, the correlation between ipsilesional TSD and lesion size was lost when photothrombosis was performed under isoflurane anesthesia (Extended Data Fig. 3g). A significant benefit of our approach is that chronically implanted optical fibers allow ischemia induction and monitoring in the absence of anesthesia or restraint, permitting evaluation of the relationship between ischemia-evoked neuronal Ca^2+^ dynamics and behavior in awake mice during the acute phase of ischemic stroke. For the remainder of the study, experiments were performed in awake (non-anesthetized) mice.

### Ischemic TSD in the hippocampus of behaving mice is linked to open field activity

In addition to amnestic deficits^65,66^, murine hyperactivity in the open field has long been reported as a behavioral proxy of lesion-induced hippocampal dysfunction^67,68^. Historically, lesions (ie. excitotoxic, mechanical, etc.) have been generated under anesthesia and animals were then tested in the following days and weeks. While there was evidence that forebrain ischemia in unanesthetized rats could trigger hyperexcitability and behavioral seizures^69,70^, it was unclear whether focal hippocampal ischemia would induce hyperactivity. We measured ischemia-evoked neuronal Ca^2+^ dynamics in freely behaving mice, allowing us to match mouse behavior to ischemia-evoked neuronal Ca^2+^ influx (i.e. SD). Mice were habituated to an open field during handling. Both sham and stroke mice gradually became less active in the open field arena over the course of the 10-minute baseline. In sham and stroke mice, we classified behavior into 4 states: sitting, ambulating, grooming, rearing (Fig. 1j) and evaluated distance travelled using DeepLabCut^71^ to identify behavior linked to the TSD in the ipsilesional hippocampus. In stroke mice, the TSD onset was identified as time of the peak derivative of the ipsilesional GCaMP6f signal (Fig. 1k). Next, the time spent in each behavioral state before and after TSD was assessed (*pre-TSD*: *-* 300 to 0 s, and *post-TSD*: 0 to + 300 s relative to the GCaMP6f peak derivative; Fig. 1*kl*). For sham mice, the average time of TSD onset in stroke mice was selected to make before/after comparisons (800 s, Extended Data Fig. 4). Following the onset of ipsilesional TSD, both female and male mice spent more time active in the open field (time spent ambulating and rearing, Fig. 1m) and travelled further (Fig. 1n-q) than sham mice, who exhibited no change in open field activity or distance travelled in an equivalent epoch (Extended Data Fig. 4). Taken together, mice increased open field activity concurrent with ipsilesional TSD.

### Female mice have more frequent contralesional SD

Following a unilateral ischemic stroke, the contralesional hemisphere is altered and plays an important role in functional recovery^72,73^. To understand the acute remote effects of unilateral hippocampal ischemia, a contralesional optical fiber was implanted above the homotopic hippocampus, naïve to photothrombotic or sham illumination (see Fig. 1c). During baseline, the frequency of basal GCaMP6f events was not significantly different in opposing hippocampi, nor between sham and stroke mice (Extended Data Fig. 5). Following ipsilesional TSD, large (∼100 fold of basal GCaMP6f dynamics) and slow contralesional neuronal Ca^2+^ transients (GCaMP6f) were observed (Fig. 2a-c) at an average of 382.6 (± 45.71) and 755.0 (± 105.6) seconds after the ipsilesional TSD for females and males, respectively (Fig. 2, Extended Data Fig. 6). These contralesional GCaMP6f transients were more frequent in females (63%) than males (25%, Fig. 2c, Extended Data Fig. 6) with no sex differences in amplitude (ΔF/F) when they did occur (Fig. 2d). We suspected that these massive fluctuations in GCaMP6f were the Ca^2+^ component of SD for several reasons^29^. First, the large amplitude of these GCaMP6f transients could only be attributed to near pan-neuronal depolarization in the fiberoptic fluorescence collection volume. Second, the period of the GCaMP6f transient (distance between start and end, female: 56.2 ± 2.4 s, male: 55.6 ± 6.4 s, Extended Data Fig. 6) approximated SD propagation recorded using neuronal Ca^2+^ biosensors through other optical methods^31,48^. Lastly, both the basal GCaMP6f level and the frequency of basal GCaMP6f events (>4 standard deviations ΔF/F) were markedly reduced for several minutes following these contralesional GCaMP6f waves (Fig. 2e-g), suggesting an accompanying depression. The reduction in basal GCaMP6f events was negatively correlated to the amplitude of the massive GCaMP6f transient (Fig. 2h). Thus, from here forth, we will refer to these large contralesional GCaMP6f transients as contralesional SD. The presence and amplitude of contralesional SD had no significant relationship to the amplitude of ipsilesional TSD (Extended Data Fig. 6).

**Figure 2.**
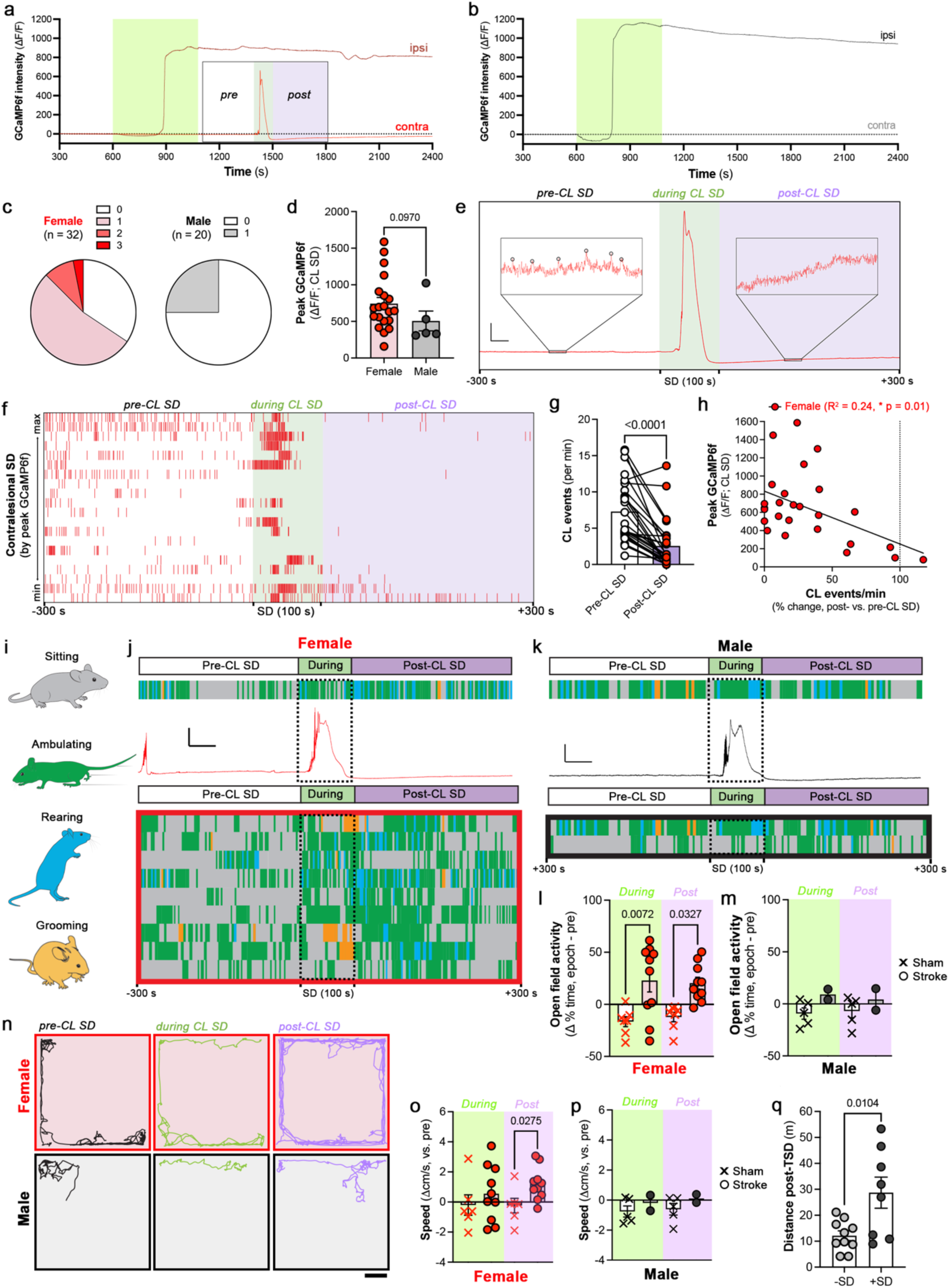
Female mice have more frequent contralesional SD. Representative IL (dark) and CL (light) GCaMP6f (ΔF/F) in **a** female (red) and **b** male (black) mice experiencing unilateral hippocampal photothrombosis. **c** CL SDs post-IL TSD per mouse by sex. **d** No sex differences in CL SD amplitude (ΔF/F). **e** Representative female CL SD highlights reduced basal Ca^2+^ events following the large GCaMP6f transient. Scale bar: 200 arb. units, 30 s. Basal Ca^2+^ events (circled, peaks > 4 standard deviations of mean, separated by > 0.5 s;) are reduced post-CL SD (frames: 15 arb. units, 30 s). **f** Raster plot of peri-CL SD epochs (one per row) represents that **g** CL Ca^2+^ events decrease after CL SD. **h** CL SD (ΔF/F) negatively correlates with the change (%) in Ca^2+^ events/min post-SD. **i** Behavior state (sitting – grey, ambulating – green, rearing – blue, and grooming – orange) during peri-SD epoch (SD – 1 min window, ± 5 min). Behavior state (Gantt plot) aligned to representative **j** female (red) and **k** male CL SDs (ΔF/F, black). Scale bar: 100 arb. units, 50 s. Gantt behavioral plots for each **j** female (red) and **k** male (black) centered on CL SD (one mouse per row). **l** Females increase time active (% of epoch ambulating or rearing) during and after CL SD. **m** Male activity is unaffected by CL SD. **n** Locomotion traces before (*pre-CL SD), during* and after (*post-CL SD)* CL SD in a representative female (red) and male (black). **o** Average speed is unchanged during yet increases following CL SD in females. **p** Average speed is unchanged during/after CL SD in males. **q** CL SD in stroke mice increases distance travelled post-TSD. Additional statistical information is available in Supplementary Information. Source data are provided in the Source Data file.

### Female mice increase time spent active during and after contralesional SD

We next asked if SD propagation through perfused tissue, as in the contralesional hippocampus, drove similar changes in behavior as ischemic TSD in the context of unilateral hippocampal photothrombosis. In stroke mice, we classified behavior before, during, and after contralesional SD (*pre-SD*: *-*350 to -50 s relative to, *during*: 100 s surrounding, and *post-SD*: +50 to +350 s relative to center of SD, respectively; Fig. 2i-k). Females further increased open field activity (i.e. above the previous ischemia-evoked TSD increase) both during and after contralesional SD (Fig. 2l). Within the limitations of a very low event rate of contralesional SD in males, this increase was not observed in males (Fig. 2m). While average speed was not significantly different during contralesional SD, female mice moved faster in the *post-CL* epoch (Fig. 2no). Males showed no change in average speed in relation to contralesional SD (Fig. 2p). The presence of a contralesional SD significantly increased the distance travelled in the 20 min following ipsilesional TSD (Fig. 2q). In sham mice, no change in open field activity occurred around the average time of contralesional SD in stroke mice (1400 s, Extended Data Fig. 7). Together, more frequent contralesional SD in females was accompanied by a further increase in open field activity relative to sham conditions or stroke mice absent of contralesional SD.

### Contralesional SD abrogates memory impairment following hippocampal stroke

Given the acute behavioral changes observed during TSD and contralesional SD, we next asked if ipsilesional TSD and contralesional SD contributed to hippocampal impairments following stroke. Mice with hippocampal dam age (via excitotoxic or mechanical lesion) show impaired context-dependent memory^65,66^; therefore, we used contextual fear conditioning, a hippocampus-dependent task^74–76^ surrounding unilateral hippocampal photothrombosis (*peri-stroke contextual fear conditioning*, Fig. 3a) to assess the impact on hippocampal function (Fig. 3a-b). Mice were subjected to 3-foot shocks in a novel context prior to unilateral hippocampal photothrombosis or sham illumination (∼10 min; Fig. 3a). Sham and stroke mice were then returned to the same context 24 h after illumination. Female and male sham mice increased time freezing on exposure to the context on day 2, relative to the epoch preceding foot shocks in the prior session (Fig. 3a), demonstrating contextual fear conditioning. Meanwhile, mice that received unilateral hippocampal photothrombosis showed no increase in freezing behavior when returned to the fear conditioning box, indicating impaired contextual fear conditioning. The presence of contralesional SD had no appreciable effect on peri-stroke contextual fear conditioning. Freezing in stroke mice during peri-stroke contextual fear conditioning was negatively correlated to the amplitude of ipsilesional TSD and unimpacted by contralesional SD, suggesting neuronal Ca^2+^ influx in ischemic core was related to this hippocampal dysfunction (Fig. 3b).

**Figure 3.**
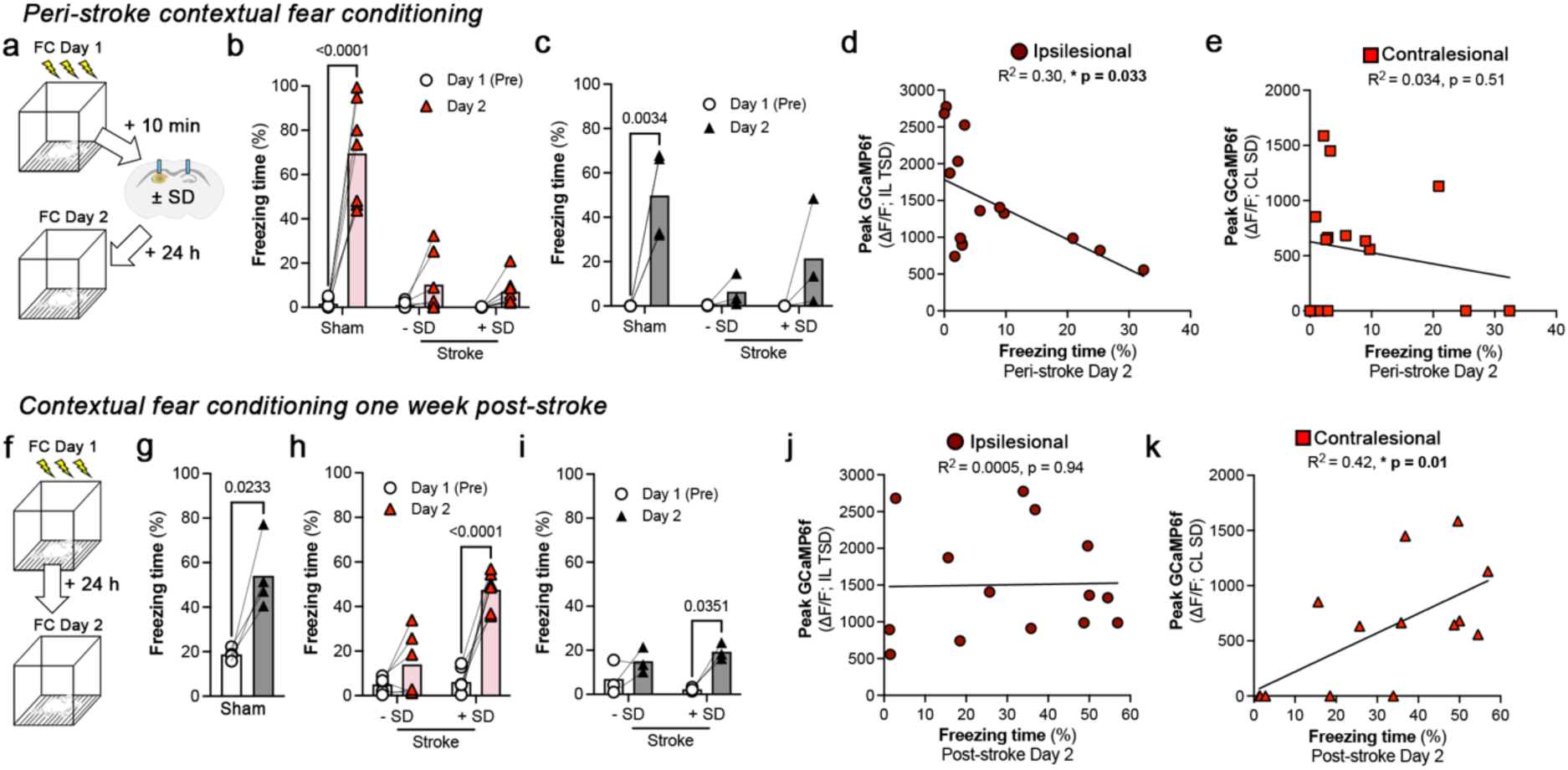
Contralesional SD is associated with improved contextual fear conditioning following hippocampal stroke. **a** Peri-stroke contextual fear conditioning. Day 1 (10 min pre-sham/stroke): 3 evenly spaced foot shocks (FS, Context 1, 7 min) followed by sham or photothrombosis. Day 2 (24 h post-stroke): mice return to Context 1 without FS (7 min). Unilateral hippocampal photothrombosis disrupts contextual fear conditioning in **b** female and **c** male mice. Time spent freezing (%) in Day 1 (pre-shock, circle) and Day 2 (triangle) for sham and stroke (± SD) mice. **d** IL TSD (***i***, ΔF/F) negatively correlates with freezing on Day 2 in females. **e** No correlation exists between CL SD and freezing on Day 2 in females. **f** Post-stroke contextual fear conditioning one week after sham/stroke. Day 1: 3 evenly spaced FS (Context 1, 7 min). Day 2: mice return to Context 1 without FS (7 min). Fear conditioning is successful in **g** sham mice one week following illumination (561 nm). Freezing time (%) increases in Day 2 (triangles) vs. Day 1 (circles) prior to FS. **h** CL SD in females recovers contextual fear conditioning to sham levels one-week post-stroke (freezing time, %). **i** CL SD in males improves contextual fear conditioning (Freezing time, %, Day 2). **j** IL TSD (***i***, ΔF/F) is not correlated with freezing on Day 2 in females (one week-post stroke). **k** CL SD (ΔF/F) correlates with increased freezing (Day 2) in females (one-week post-stroke). Additional statistical information is available in Supplementary Information. Source data are provided in the Source Data file.

One week after sham or stroke, mice were fear conditioned to assess the recovery of hippocampal function (*post stroke contextual fear conditioning*, Fig. 3c-d), with translational relevance to anterograde amnesia reported in humans^11^. Mice were placed in the fear conditioning box and 3-foot shocks were delivered over a period of 7 min. When returned, 24 h later, to the identical context for a 7 min period in the absence of foot shocks, sham mice increased time spent freezing (Fig. 3c). In stroke mice, only those that had contralesional SD following unilateral hippocampal photothrombosis, 1 week earlier, showed evidence of contextual fear conditioning. This effect was strongest in females, who recovered to sham levels (Fig. 3c). Freezing during this task was correlated to the amplitude of contralesional SD (Fig. 3d) and had no statistical relationship with the amplitude of ipsilesional TSD. Finally, freezing was specific to the fear conditioned context, as neither sham nor stroke mice increased freezing in a novel context absent of foot shock (Extended Data Fig. 8). Together, this data suggests that contralesional SD during the acute phase of stroke protected post-ischemic hippocampal function after loss of the ipsilesional hippocampus due to stroke.

### Contralesional SD is preceded by electrographic seizure

In addition to the prominent role of SD in ischemic tissue, ischemia can evoke several other electrophysiological phenomena including epileptic seizures^77^. In humans, electrographic seizures are apparent in acute continuous electroencephalogram (EEG) recordings during the early phase of ischemic stroke in up to 44% of patients, depending on lesion location and patient demographics^78–80^. Untreated seizure after stroke, can trigger post-stroke epilepsy^81^ and is associated with neurological deterioration^78^. Seizure and SD can co-occur in both rodents and humans^49–54,82–84^. While a complex pattern between SDs and electrographic seizure exists in patients receiving neurocritical care, In pharmacologically induced preclinical seizure models^83^, focal seizures can trigger a transient SD, which, after a phase of activity depression, can at least temporarily restore normal spontaneous activity^50,85^. To determine if endogenous hippocampal SD can terminate ischemia-evoked contralesional seizures as a potentially protective mechanism, we implanted a bipolar depth electrode ∼ 2 mm posterior to the contralesional fiber in female and male mice and performed unilateral hippocampal photothrombosis (Fig 4a). Following TSD in the ipsilesional hippocampus, epileptiform activity occurred in the contralesional hippocampal AC-coupled local field potential (LFP) recordings. Unilateral ischemia in both sexes elicited electrographic seizure in the contralesional hippocampus with or without coincident changes in neuronal GCaMP6f (see Fig 4). While electrographic seizures could occur independent of large fluctuations in GCaMP6f (neuronal Ca^2+^), contralesional SDs were always preceded by epileptiform activity (Fig. 4a-f). Here, electrographic seizures persisted for > 10 s prior to SD onset, were of higher power relative to non-SD-associated seizure and were followed by a transient reduction of power indicative of an accompanying spreading depression (Fig 4def). In female and male stroke mice without contralesional SD, seizures were of lower overall power and typically lasted for < 10 s without consistent changes to GCaMP6f intensity (Fig. 4f-j). Seizures directly preceding SD were longer than those that did not precede SD (Fig 4k), yet the cumulative duration of electrographic seizures (between ipsilesional TSD and the end of recording) was not significantly different between sexes (Fig 4kl). As seizures were equally present in stroke mice without contralesional SD, seizure termination alone was not a sufficient explanation for the protective effect of SD.

**Figure 4.**
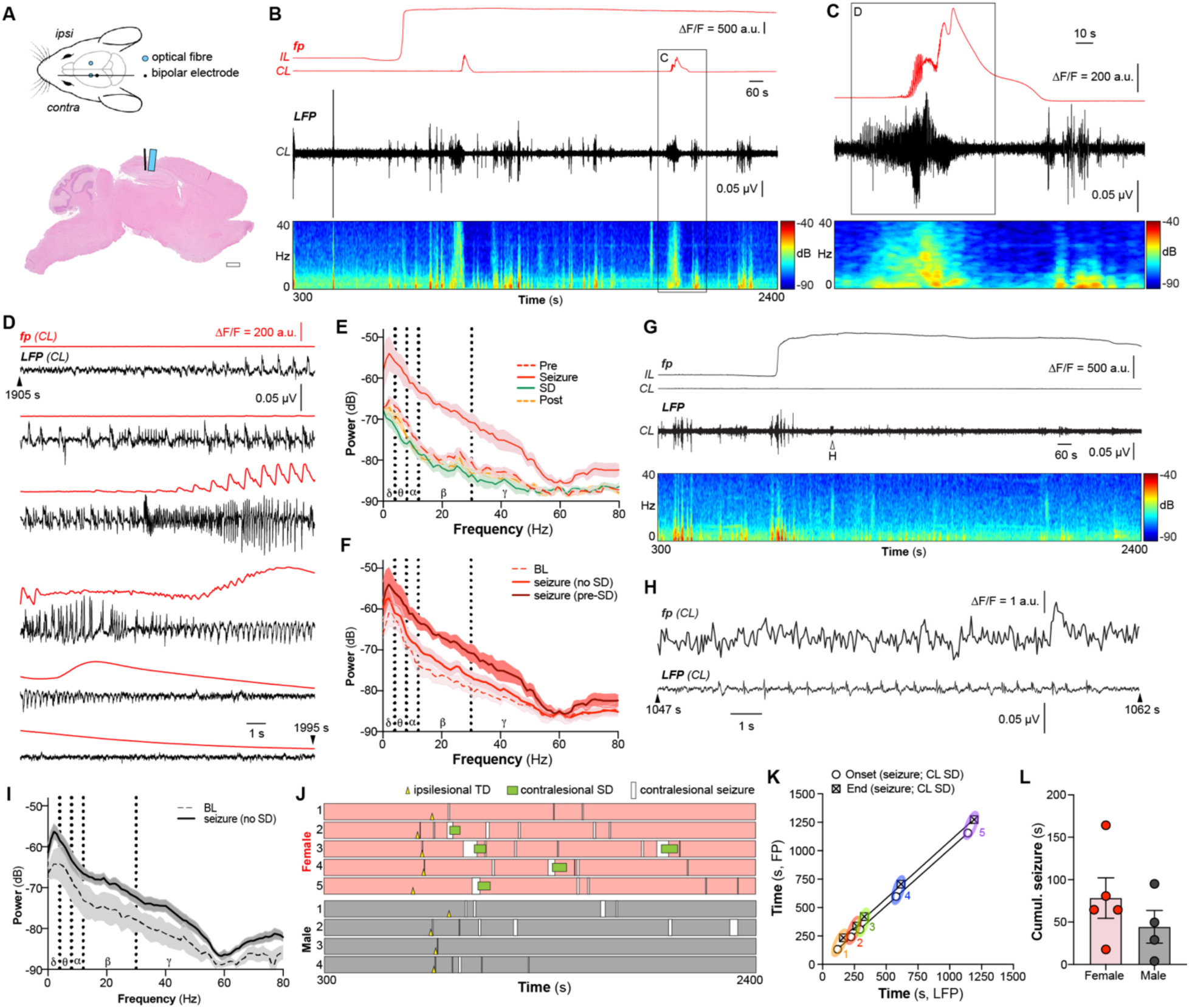
Electrographic contralesional seizure precedes SD. **a** CA1-targeted bilateral fiberoptic cannulas (blue) and CL bipolar electrode (black). Representative hematoxylin and eosin-stained sagittal section at CL fiber and electrode implant locus one-week post-stroke. Scale bar, 2 mm. **b** Example IL and CL GCaMP6f (ΔF/F, red) paired with CL LFP (black trace) and time-varying frequency-power spectrogram in an awake, behaving female mouse. Photothrombotic illumination (561 nm: 6 mW, 8 min). **c** 180 s surrounding a representative CL seizure (black, LFP) preceding an SD (red, GCaMP6f). **d** Fig6c inset highlights seizure (LFP) preceding CL SD (GCaMP6f) followed by depression in LFP power. **e** Power spectral density (20 s epochs) of contralesional LFP (pre-SD: 300 s preceding the peak of CL SD; seizure: during the seizure that directly precedes SD; SD: during the peak of CL SD; and 300 s after the peak of CL SD) in females. **f** Power spectral density of 60 s epochs encapsulating baseline (400-460 s), seizures that directly precede SD (pre-SD), or 20 s epoch surrounding seizures without SD (no SD) in females. **g** Example IL and CL GCaMP6f (ΔF/F, black) paired with CL LFP (black) and time-varying frequency-power-time spectrogram in an awake, behaving male mouse. **h** Expanded *Fig6e (white arrow)* highlights epileptiform activity (LFP) in 15 s window. **i** Power spectral density of contralesional LFP in male stroke mouse for 60 s epochs encapsulating baseline (400-460 s) or 20 s epoch surrounding seizures that occur independent of SD (no SD) in males. **j** Schematic overlay of CL recordings (LFP/FP) in female (red) and male (black) mice, highlighting seizure incidence (white box) in relation to IL TSD (yellow triangle) and CL SD (green box). **k** Onset (empty circle) and end (hatched box) of CL seizure (LFP, x-axis) and SD (GCaMP6f FP, y-axis) in the hippocampus following IL TSD is correlated. **l** Cumulative duration of CL seizure following IL TSD. Additional statistical information is available in Supplementary Information. Source data are provided in the Source Data file.

### Contralesional SD arises from secondary contralesional seizure

The ubiquity of electrographic seizure preceding contralesional SD (Fig. 4) led us to explore if seizure, rather than SD, was propagating outward from the ischemic tissue into the opposing hippocampus. We performed bilateral photometry paired with bilateral hippocampal LFP (Fig. 5a) during unilateral hippocampal photothrombosis in female mice. Bipolar depth electrodes were implanted at a posterior tilt, so that the tip of each electrode was ∼2 mm posterior to its ipsilesional optical fiber. Seizures are identified in an LFP recording as increased peak amplitude and overall signal power, with reduced entropy, and increased rhythmicity. Using a custom-written MATLAB script, we identified ‘suprathreshold peaks’ in the LFP signal that were ≥ 4 standard deviations from the mean of noise-free baseline LFP. We then determined if the subsequent LFP signal had evidence of hyper-synchronicity and rhythmic spiking, indicative of an electrographic seizure. In each of the bilateral LFP recording where contralesional hippocampal seizure was immediately followed by an SD, suprathreshold rhythmic spiking appeared first in the contralesional LFP before appearing in the ipsilesional LFP (Fig. 5b). In three of the four mice where contralesional seizures were observed and SD followed, contralesional seizure preceded the onset of ipsilesional hippocampal seizure (by 2.81 ± 0.76 s, Fig. 5b) and, in one mouse, seizure was restricted to the contralesional hippocampus. Over a 60 s period encapsulating the pre-SD seizure, LFP power (via short-time Fourier transform) was binned in overlapping windows (1 s bins, 512 points in fast Fourier transform) and plotted over time. Next, each plot was normalized to the maximum amplitude of the power during the contralesional seizure and aligned to the initial suprathreshold peak in the contralesional LFP. LFP power (0 - 80 Hz, Fig. 5b-d) increased to a greater degree in the contralesional hippocampus prior to changes in LFP from ipsilesional hippocampus, indicating the generation of a secondary seizure focus that had the potential to weakly propagate to the ipsilesional hemisphere. To rule out that interhemispheric SD propagation was occurring via the superficial cortex, we used mesoscale cortical imaging^86,87^ of Thy1-GCaMP6f mice paired with bilateral fiber photometry (Fig. 5e). Optical fibers were implanted at a posterior and lateral tilt to optimize the mesoscale field of view and achieve simultaneous fiber photometry from hippocampi and fluorescence imaging from a large expanse of dorsal neocortex. This preparation required head-fixation to optimize the stability of the imaging plane. Structure-associated ROIs (11 px^2^, 35µm/pixel) on the superficial cortex were identified based on manual alignment to the Allen Brain Atlas. Following unilateral hippocampal photothrombosis, no appreciable SD was observed in the superficial cortex of either hemisphere (Fig. 5g), despite SD appearing in the contralesional hippocampus (n = 5, 3 with hippocampal SD, 2 without hippocampal SD, Fig. 5g). Slowly propagating GCaMP6f transients were visible in the neocortical structures neighboring the ischemic hippocampus following unilateral hippocampal photothrombosis (Fig. 5g, light lines); however, they were much smaller in amplitude (ΔF/F ≈ 10 a.u.) relative to those that appeared following photothrombosis in neocortical tissue (ΔF/F ≈ 200 a.u., Extended Data Fig. 9). Rhythmic seizure-like GCaMP6f dynamics were observed in contralesional mesoscale recordings from nearby cortical brain areas. This suggests that the seizures preceding hippocampal SD could be propagating into neocortex. Given that SD involves pan-neuronal depolarization^20^, the accompanying slow and small amplitude GCaMP6f transients (ΔF/F ≈ 10 a.u.) in contralesional neocortical structures were not indicative of SD entering these structures (Extended Data Fig. 10). Here, the increase in GCaMP6f was between 5-10% of what was observed during cortical SD in these same structures after cortical photothrombosis (Extended Data Fig. 10). In control experiments, a larger unilateral infarction that affected both the hippocampus and superficial cortex did trigger a cortical SD; however, in this case, SDs neither crossed the midline into the contralesional neocortex nor evoked a contralesional SD in the hippocampus (Extended Data Fig. 9).

**Figure 5.**
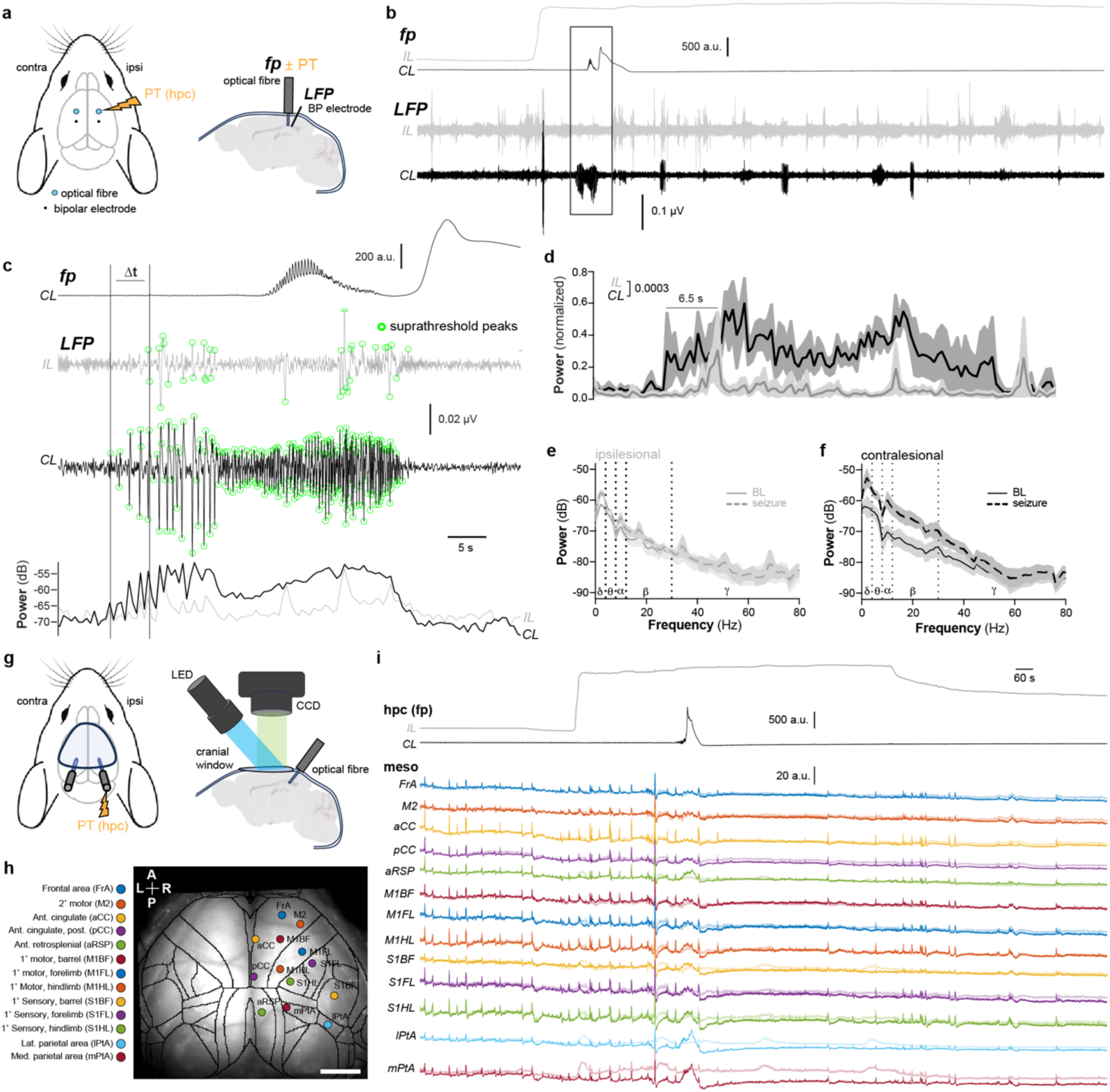
Contralesional SD arises from secondary contralesional seizure. **a** Bipolar electrodes implanted posterior to fiberoptic cannulas at a 10° posterior tilt. **b** IL (grey) and CL (black) GCaMP6f (ΔF/F) paired with respective LFP in an awake, behaving female mouse. **c** 60 s window preceding and overlapping with CL SD. Suprathreshold peaks (> 4 standard deviations, LFP, green circles) indicating contralesional seizure preceded onset of suprathreshold ipsilesional peaks. Increase in contralesional LFP power (0-80 Hz, low pass (20 Hz) filtered) precedes ipsilesional LFP power increase. **d** Average power of bilateral LFP aligned to initial suprathreshold peak of CL LFP (CL peak at 10 s, total 60 s). Power spectral density for **e** ipsilesional (light) and **f** contralesional LFP (dark) from 60 s of noise-free baseline LFP (basal) relative to 60 s overlapping seizure (seizure). **g** Mesoscale imaging window and bilateral fiberoptic implants targeting dorsal hippocampi for simultaneous mesoscale GCaMP6f imaging in the superficial neocortex with unilateral hippocampal photothrombosis and bilateral photometry. **h** Dorsal neocortex during mesoscale image acquisition highlighting region-specific ROIs. **i** Bilateral hippocampal photometry and structure/ROI-based (11 px^2^, 35 µm/px) mesoscale imaging traces for ipsilesional (light, bold) and contralesional (dark) neocortical structures following unilateral hippocampal photothrombosis with subsequent CL SD highlight the absence of interhemispheric cortical SD propagation (n = 5, 3 with CL hippocampal SD, 2 without CL hippocampal SD). Additional statistical information found in Supplementary Information. Source data are provided in the Source Data file.

### Blunting contralesional SD worsens post-stroke contextual fear conditioning

We next aimed to specifically disrupt contralesional SD, while leaving ipsilesional peri-infarct depolarization intact, to understand its potential role in neuroprotection. NMDA receptors contribute to SD initiation and propagation^50,88–91^ with their precise role depending on stimulus, tissue health, and context^20,64^. There is evidence that low dose (0.5 mg/kg) of the pore-blocking, uncompetitive NMDAR antagonist MK801 prevents SD propagation into healthy tissue without affecting SD in severely ischemic tissue^92^. MK-801 was also selected given its ability to blunt SD without preventing seizure^50^, which further allowed us to isolate SD-dependent outputs. MK-801 was delivered i.p. (0.5 mg/kg, Fig. 6) immediately prior to baseline and 10 min prior to sham illumination or photothrombosis to assess whether contralesional SD was necessary for intact contextual fear conditioning one-week post stroke. MK-801 reduced the amplitude of ipsilesional TSD by 35.3 ± 15.4% relative to vehicle-treated mice (Fig 6a-c). MK-801, had an outsized effect on contralesional SD, reducing their amplitude by 66.0 ± 9.4% relative to vehicle (Fig. 6d). Seizures preceding SD in MK-801-treated mice had larger amplitude (Fig. 6e) and increased power, relative to vehicle. Following MK-801 treatment, the electrical depression after contralesional SD was either absent or abbreviated (Fig. 6b,f). Further, MK-801 increased the cumulative duration or contralesional seizures in the recording period after ipsilesional TSD relative to vehicle (Fig. 6gh, Extended Data Fig. 11). In vehicle-treated mice, SD improved contextual fear conditioning one week after stroke relative to mice without SD (Fig. 6j), consistent with results from naïve mice (Fig. 3c). In the same assay, sham mice treated with MK-801 immediately prior to photo-illumination demonstrated contextual fear conditioning (Fig. 6j). MK-801-treated stroke mice, in which SDs were blunted, did not demonstrate contextual fear conditioning when trained one-week post-stroke. This provides evidence that intact contralesional SD is necessary to preserve non-infarcted hippocampal function following unilateral hippocampal stroke.

**Figure 6.**
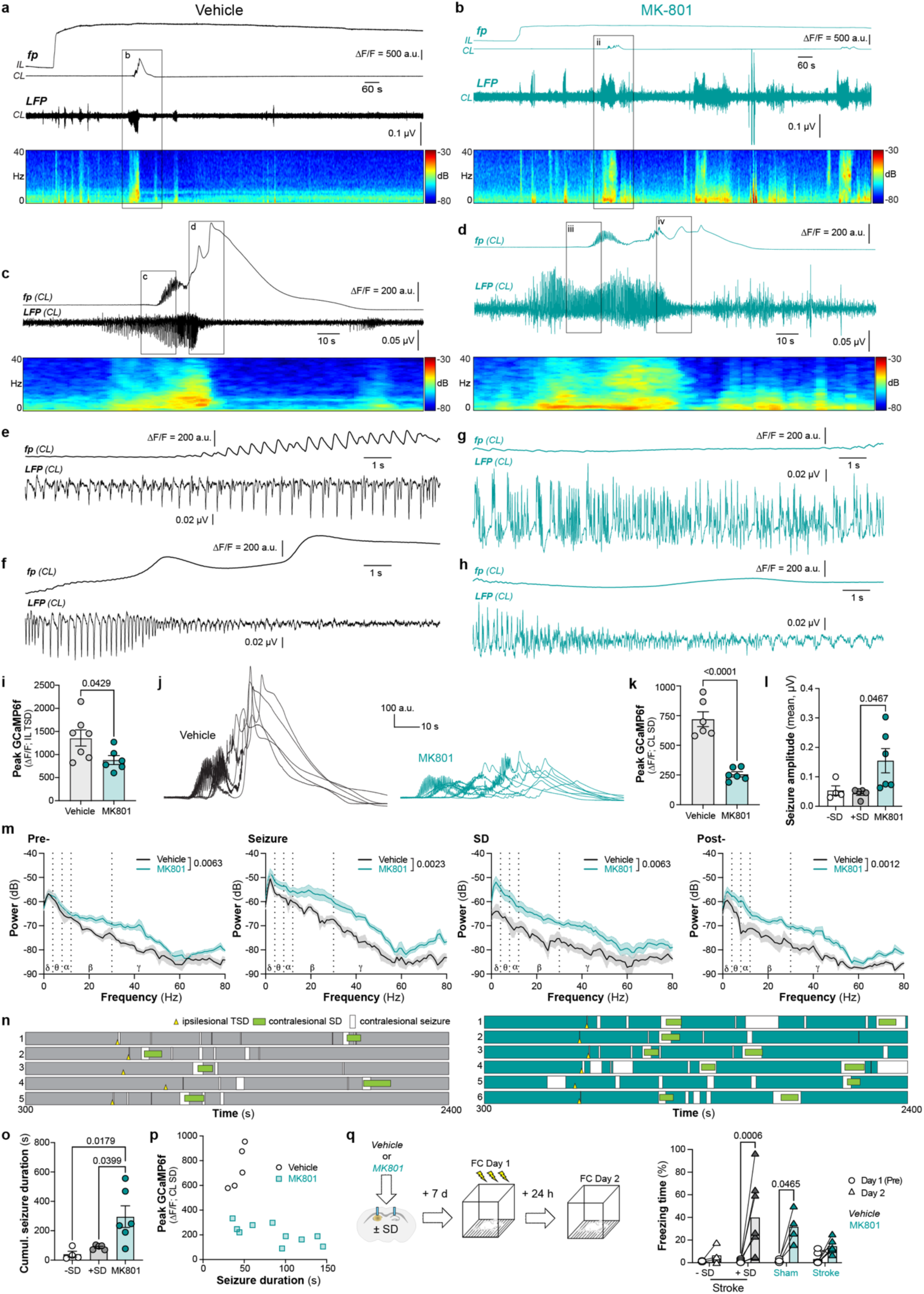
Contralesional SD arises from secondary contralesional seizure. **a** Bipolar electrodes implanted posterior to fiberoptic cannulas at a 10° posterior tilt. **b** Representative IL (grey) and CL (black) GCaMP6f (ΔF/F) paired with respective LFP in an awake, behaving female mouse. **c** 60 s window preceding and overlapping with CL SD. Suprathreshold peaks (> 4 standard deviations, LFP, green circles) indicating contralesional seizure preceded onset of suprathreshold ipsilesional peaks. Increase in contralesional LFP power (0-80 Hz, low pass (20 Hz) filtered) precedes ipsilesional LFP power increase. **d** Average power of bilateral LFP aligned to initial suprathreshold peak of CL LFP (CL peak at 10 s, total 60 s). Power spectral density for **e** ipsilesional (light) and **f** contralesional LFP (dark) from 60 s of noise-free baseline LFP (basal) relative to 60 s overlapping seizure (seizure). **g** Mesoscale imaging window and bilateral fiberoptic implants targeting dorsal hippocampi, allowing simultaneous mesoscale GCaMP6f imaging in the superficial neocortex with unilateral hippocampal photothrombosis and bilateral photometry. **h** Dorsal neocortex during mesoscale image acquisition highlighting region-specific ROIs. **i** Representative bilateral hippocampal photometry and structure/ROI-based (11 px^2^, 35 µm/px) mesoscale imaging traces for ipsilesional (light, bold) and contralesional (dark) neocortical structures following unilateral hippocampal photothrombosis with subsequent CL SD highlight the absence of interhemispheric cortical SD propagation (n = 5, 3 with CL hippocampal SD, 2 without CL hippocampal SD). Additional statistical information found in Supplementary Information. Source data are provided in the Source Data file.

## Discussion

We present a model for both inducing ischemic stroke and recording subsequent neuronal Ca^2+^ dysregulation in awake, freely behaving mice. Symmetric bilateral fiberoptic cannulas targeting the CA1 in Thy1-GCaMP6f mice revealed ipsilesional terminal SD and associated behavior that was similar between sexes, yet revealed a strong sex difference in contralesional SD. These waves immediately followed seizures, increased open field activity (ambulating and rearing) upon arrival, and had longitudinal protective effects on contextual fear conditioning during stroke recovery. Here, we have demonstrated key differences between acute and long-term effects of hippocampal stroke relative to other brain regions. This is important because it supports the clinical observation that stroke in PCA-fed territory has consequences on behavior, memory and long-term cognition that require unique standards of care^11^.

Large vessel occlusions in MCA-fed territory account for approximately half of ischemic strokes in humans^2^. Several preclinical models exist for focal ischemic stroke in the superficial cortex, including the use of photothrombosis^48,60,93^ and aggregation of magnetic nanoparticles^94^. Despite the remainder of strokes occurring in territories not supplied by MCA, considerably less attention has been given to developing models for stroke in deeper brain structures. There are several models where stroke encroaches on or is targeted to deeper structures, each with unique benefits and limitations. Intraluminal middle cerebral artery occlusion (MCAO) induces a large lesion that affects the striatum and can spread into superficial neocortex, while also reaching other subcortical regions when occluded for prolonged periods^3^, however MCAO requires anesthesia and is highly variable. Injection of thromboemboli or microspheres can induce subcortical stroke^95^, but also creates unpredictable infarcts distributed diffusely throughout the brain and also requires anesthesia. Importantly, regional specificity of micro-infarctions is dictated by methodology and size of occluding material^95,96^. Optical fibers have been used for photothrombosis beneath the superficial cortex^97–99^, yet none of these studies paired photothrombosis with an acute readout of neuronal depolarization (ie. GCaMP6f photometry) or used awake unanesthetized mice, as in our study. Anesthesia is protective against stroke and inhibits SD^60–62^. Not only does anesthesia eliminate the ability to understand mouse behavior associated with ischemia, isoflurane acts competitively on the glycine site of NMDARs^63^ and strongly inhibits SD^61,62^, which can mask effects of other NMDAR antagonists. This is particularly important given that NMDAR antagonism has been the most popular and potent pharmacological approach for inhibiting SD^64^. While focal stroke can be triggered in the superficial cortex of awake head-fixed mice^48,94^, these approaches also restrict behavior evaluation during ischemia induction.

To better understand stroke in deeper brain structures, we began by targeting the hippocampus, a brain structure involved in episodic and contextual memory and spatial navigation across species^5,6^ with well-defined behavioral assays to specifically target hippocampus-dependent tasks^5^, allowing clear demonstration of the utility of our model in deeper brain structures. Strokes in the posterior cerebral artery (PCA), the vessel that supplies the hippocampus, account for about one-third of all non-middle cerebral artery (MCA) single vessel strokes. Clinical presentations of hippocampal stroke include confusion and disorientation, which can resolve in the weeks or months following stroke^11–13^, along with persistent memory deficits^13,100^. In humans, unilateral hippocampal infarcts can induce verbal or non-verbal long-term episodic memory deficits depending on left or right lesion location, respectively^11^. In our study, increased activity in a familiar environment occurred concurrent with the onset of TSD in the ipsilesional hippocampus (Fig. 1) and further increased with SD in the contralesional hippocampus (Fig. 2). While hippocampal strokes in humans are rare, a recent report described complementary findings in another disease model (viral encephalitis) and in humans with temporal lobe epilepsy^101^. They describe *post-ictal wandering* triggered by hippocampal seizure-induced SD^101^, highlighting that this phenomena may occur beyond the stroke brain. Here, we identify hyperactivity in response to unilateral hippocampal ischemia and subsequent SD, which could be caused by (1) impaired spatial memory, reintroducing a “novel” environment, and slowing habituation^102^ or (2) a reduction of anxiety, given that inhibiting ventral hippocampus function can be anxiolytic^103^, or a combination of both factors. The hippocampus interacts with several brain structures involved in modulating the stress response^104^, including integrating ‘cue’ and ‘context’ information. The hippocampus can also regulate spatial memory^105^ and indirectly influence running speed^106,107^, both of which could impact navigation. There is evidence for delayed spatial memory impairments in complex environments following dorsal hippocampal infarcts in rats^108^, however these assessments were done following recovery from a stroke under anesthesia. The underlying mechanism and motivation for hippocampal SD-induced hyperactivity should be the focus of future research.

Symptomology of stroke is linked to the affected brain region. Here, we observed an increase in ambulating and rearing in the open field linked to hippocampal SD. This was distinct from acute hypo-activity that was triggered by photo-induced MCAO occlusion, where mice froze during photothrombotic illumination of the MCA^60^. A caveat of this previous MCAO study was the absence of a readout of neuronal depolarization (ie. no simultaneous recording), thus photothrombotic illumination was used to approximate the onset of ischemia^60^. Together, our data and the previous study provide evidence that, like longitudinal stroke-related impairments^109^, acute behavior associated with ischemia is region-specific.

Contralesional hippocampal SD occurred in 63% of females and 25% of males after unilateral hippocampal photothrombosis (Fig. 2). There is preclinical evidence that females have a reduced threshold for SD in healthy tissue^110–113^. Sex differences in SD susceptibility have been shown in response to topical application of concentrated potassium chloride (KCl) on well-perfused neocortex^111^. In that study, female rats had increased SD frequency that was eliminated by gonadectomy. In gonadectomized females, estrogen withdrawal after prolonged daily treatment or, conversely, daily injection of progesterone increased CSD susceptibility, while this effect was not observed in males^111^. In another study, 17ß-estradiol and progesterone supplementation after ovariectomy increased CSD frequency^113^. Females had a lower SD threshold in a genetic model of familial hemiplegic migraine (FHM1) that was abolished by ovariectomy^112^. Together, these data provide evidence that gonadal hormones may regulate SD in the contralesional hippocampus after unilateral focal ischemia. The absence of a sex difference in peri-infarct TSD relative to increased contralesional SD in females suggests that the sex difference may be specific to SD initiation and propagation in ‘healthy’ tissue, or seizure conversion into SD, which is supported by the findings discussed above where sex differences in SD were explored in naïve neocortex^110–113^.

Mice experiencing unilateral hippocampal photothrombosis without contralesional SD showed impaired contextual fear conditioning both surrounding the stroke, as well as one week following stroke (Fig. 3ab), paralleling amnesia found in some hippocampal stroke patients^11^. Despite the unilateral nature of the ischemic injury, leaving the contralesional hippocampus structurally intact, freezing was absent in stroke mice during peri-stroke contextual fear conditioning (Fig. 3ab), suggesting that broader hippocampal network function and memory encoding was disrupted. Stroke mice that had a contralesional SD (in the 30 min following ipsilesional TSD) had intact contextual fear conditioning one week after stroke, suggesting a recovery of hippocampal function (Fig. 3cd). Increased freezing during the retrieval day (Day 2) correlated with contralesional SD amplitude but had no relationship to ipsilesional TSD. This suggests contralesional SD in the intact hippocampus plays a unique role in maintaining hippocampal function after a focal unilateral insult. While we provide evidence that an intact contralesional SD is necessary for contextual fear conditioning after stroke (Fig. 3 & 6), the mechanism of protection remains elusive. It is possible that ischemia-evoked contralesional SD is upregulating plasticity-inducing or neurogenic factors, as observed in other models^40–44^. Here, inhibition of the NMDAR-component of contralesional SD, and subsequent failure of contextual fear conditioning post-stroke (Fig. 6), could be due to the disruption of a protective form of synaptic plasticity. In our study, GFAP-reactive area in the contralesional hemisphere was increased following unilateral hippocampal stroke (Extended Data Fig. 2). This was consistent with a previous study that reported increased astrocyte proliferation in the contralesional hemisphere following neocortical SD induction^114^. The hippocampus is particularly benefited by the presence of SD, as KCl-evoked SD localized to the cortex induced increased adult hippocampal neurogenesis in both rats and mice^41,43^ and enhanced object location memory in the weeks following SD^41^. While one week post-stroke is insufficient time for *de novo* neurogenesis and network of newborn progenitors^115^, it is possible that SD could improve or hasten integration of existing neuronal and glial progenitors or enhance hippocampal plasticity.

Although the hippocampus is a highly epileptogenic structure, it is likely that elements of our findings will translate to other brain regions where stroke occurs. Post-stroke ictogenesis is not restricted to focal hippocampal stroke. Continuous EEG recordings reveal epileptiform activity in up to 44% of human ischemic stroke patients in acute stroke care^78^. In pre-clinical studies, post-stroke seizure in the ipsilesional hippocampus is common in photothrombotic stroke restricted to the superficial cortex, occurring in ∼ 67% of mice in the week after stroke^116^, with up to 80 seizures occurring per day. In a middle cerebral artery occlusion model, periodic epileptiform discharges were observed in simultaneous cortical EEG recordings of ∼ 80% of rats^117^ (however, see ^14^).

In our study, seizure occurred in the contralesional hippocampal LFP of all mice following unilateral hippocampal photothrombosis, yet not all seizures led to SD (Fig. 4). However, in all cases of contralesional SD, epileptiform activity preceded the GCaMP6f response (Fig. 4, n = 5; Figure 5, n = 4). The most common location for a secondary seizure is the homotopic contralateral region to a primary injury, forming a so-called ‘mirror focus’^118^. Mirror foci have been described in mammalian hippocampi after unilateral excitotoxic lesion or seizure^119–123^ and can appear within 10 min of an initial injury^123,124^. The hippocampus forms strong commissural connections with its homotopic structure in the contralateral hemisphere^125–127^. In an *in vitro* preparation consisting of physically isolated bilateral hippocampi and their commissural fibers alone, it was possible to generate a rapid onset mirror focus in a naïve hippocampus following a unilateral lesion^123^. Following unilateral kainic acid (KA) injection in rats, a spontaneous rapid onset mirror focus (< 10 min post-injection) formed in the contralesional hippocampus and preceded ipsilesional seizure activity in 73% of the recordings (by an average of ∼ 5 s)^120^. Contralesional mirror-focus development was AMPAR-dependent, as AMPAR antagonism biased seizure presentation to the ipsilesional hippocampus, while kainate receptor antagonism was ineffective at switching onset location^120^. The role of AMPARs in initiating seizure activity at mirror foci was supported by an additional study in *ex vivo* guinea pig brain^124^. This could be explained by enhanced excitatory drive initiated by an ischemic TSD in the ipsilesional hippocampus that could propagate through commissural fibers to the contralesional hippocampus. There are several potentially co-operative mechanisms through which unilateral hippocampal ischemia could trigger acute seizure activity in the contralesional hemisphere. Extracellular [K^+^] is increased in ischemic tissue^128^, with activation of neuronal K^+^ channels, and disrupted Na^+^/K^+^ pump activity, ultimately lowering the threshold for seizure and SD. Neurotransmitter release is also enhanced in ischemic tissue, further increasing seizure susceptibility^129^. In peri-infarct tissue, blood brain barrier dysfunction causes albumin to leak into the brain parenchyma, which can also activate astrocytic transforming growth receptor beta receptor 2 (TGFßR2) signaling, altering the astrocyte transcriptome to disrupt local [K^+^] and glutamate homeostasis and enhance seizure activity and epileptogenesis^116,130,131^. However, the latter mechanism is likely not feasible given the rapid onset of seizure after photothrombosis in our study (Fig. 4). In mice and rats, CA3 pyramidal cells have commissural axons that form predominantly excitatory synapses with both pyramidal cells and interneurons of contralateral CA3 and CA1^125–127^. Here, AMPAR-mediated activation of resident inhibitory interneurons can synchronize network oscillations and initiate epileptiform activity in the contralesional hippocampus^132^.

NMDAR antagonism of the contralesional SD (via MK-801) impaired the functional recovery of the remaining hippocampus following unilateral hippocampal stroke (Fig. 6). NMDAR antagonists have a dose-dependent and context-specific inhibitory effect on SD^92,133,134^. For instance, at higher doses (> 3 mg/kg), MK-801 consistently and robustly reduces ischemic lesion volume^135^. At lower doses (0.5 - 1 mg/kg), the neuroprotective effect of MK-801 against ischemic stroke is contested (^136,137^ yet see ^138,139^). During photothrombosis in rats, 0.5 mg/kg MK-801 effectively blocked SD propagation outward from penumbra into sufficiently perfused tissue, but was ineffective at preventing TSD in core/penumbra^92^. Therefore, it is not surprising that disruption of contralesional SD with low dose MK-801 led to worse performance during contextual fear conditioning one-week post stroke (Fig. 6). This is not a direct effect of MK801 on fear conditioning, as MK801 was given in a single injection one-week prior to the acquisition round of post-stroke contextual fear conditioning. Specific inhibition of SD in adequately perfused tissue was our justification for using 0.5 mg/kg MK-801; however, low dose MK-801 impaired locomotion of mice in our study. Ketamine, a less potent uncompetitive antagonist of NMDARs that shares a binding site with MK-801^140^, can also suppress SDs^14^. Ketamine blocks SD propagation, then initiation, in a dose-dependent manner^134^. Importantly, subanesthetic doses of ketamine is used in humans for treatment of depression^141^ among other neurological disorders^142^. Ketamine inhibits SD at anesthetic doses in rodents^133^ and humans ^143–146^ Subanesthetic ketamine can reduce the length of the DC shift and the period where evoked potentials are suppressed during SD in an *ex vivo* hippocampal slice preparation^133^. Neuronal Ca^2+^ load could be reduced by subanesthetic ketamine *in vivo* at concentrations that did not inhibit SD^147^. In future studies, subanesthetic ketamine could be tested in sham and stroke mice, in lieu of MK-801, to tease out both the acute and long-term behavioral effects associated with contralesional SD inhibition.

NMDAR antagonists have failed in human stroke trials due to several complications, including specificity, dosing, timing of administration, and off-target cognitive effects^64^. The disruption of a protective SD following focal ischemia may be an additional reason for their failure. Importantly, NMDAR antagonists are unable to prevent the TSD that occurs during severe ischemia^148^, likely due to recruitment of other cation channels, AMPA, kainate, and metabotropic glutamate receptors, and/or non-ionotropic NMDAR signaling to pannexin-1^55,139^. While existing on a continuum, unique receptor activation and signaling during SD in severely ischemic versus well-perfused tissue could be a ‘feature, not a bug’ given that they might offer unique therapeutic targets.

Our model opens the door for using fiber photometry to study pathophysiological phenomena and offers a unique perspective on stroke biology in freely behaving mice with important flexibility to study stroke throughout the brain. We identify important and clinically relevant unique behaviors and sequalae associated with focal hippocampal stroke. Future studies could use other biosensors or explore other cell types and brain regions using this approach to better understand how brain activity in regions both proximal and distal to focal stroke are linked to peri-stroke symptomology and outcome. Together, this will help to inform stroke care in a sex and brain context-specific manner.

## Methods

### Animals

All animal care protocols were in accordance with the Canadian Council on Animal Care guidelines and approved by the University of Calgary’s Animal Care and Use Committees (protocol #AC21-0181). Both Thy1GCaMP6f (C57BL/6J-Tg(Thy1-GCaMP6f)GP5.17Dkim/J, JAX 025393)^149^ female and male mouse littermates from Thy1GCaMP6f(+/-) breeding pairs were group housed until the first procedure and then single housed on a 12 h/12 h light/dark cycle. Mice had *ad libitum* access to Purina laboratory chow and water.

### Fiberoptic and electrode implant

Thy1GCaMP6f mice (P60-120, N = 65, 32 awake bilateral females, 20 awake bilateral males, 5 awake unilateral males, 8 anesthetized unilateral males) were anesthetized using inhaled isoflurane (5% induction; 1% maintenance) for surgery, then placed in a stereotaxic apparatus. Next, the area above the skull was shaved to expose the scalp and a midline incision (∼3 cm) was made. The skull was exposed by detaching the muscle from the sagittal ridge then cleaned using 5% hydrogen peroxide. Bregma and lambda were located and equilibrated in the dorsal ventral plane. From here, mono-fiberoptic cannulas (MFC_400/430-0.48_2.5mm_ZF2.5(G)_FLT, Doric Lenses) were implanted dorsal to the CA1 region of each hippocampus (AP -2.0 mm, ML ±1.3 mm from bregma, and DV -1.0 mm from the brain surface). Where applicable, secondary burr holes were made for bipolar depth electrodes to simultaneously record local field potentials (LFPs) in either both hippocampi or contralesional hippocampus only. Electrodes were constructed from a 178 μm stainless steel wire connected to gold-plated male amphenol pins (A-M systems) and lowered at stereotaxic coordinates: (right: AP -2.30, ML +1.3 mm from bregma, DV -1.5 mm from surface; left: AP -2.30, ML -1.3mm from bregma, DV -1.5 mm from surface). Implants were fixed to the skull using serial application of Metabond and dental cement. For fiber photometry with LFP experiments, Thy1GCaMP6f mice (P60-120, N = 23, 19 awake bilateral females (5 naïve, 9 vehicle, 7 MK-801), 4 awake bilateral males) were used. Mice were allowed to recover for 10 days prior to sham illumination or photothrombosis.

### Photothrombotic stroke & fiber photometry in awake and freely behaving mice

For focal ischemia in the hippocampus (photothrombosis^47^) in awake and behaving mice, vehicle (PBS pH 7.2, 0 MgCl_2_, 0 CaCl_2_; Invitrogen Cat. 20012050) or Rose Bengal (1% w/v in PBS; Sigma Canada R3877-5G) were administered via intraperitoneal injection (0.1 mL per 10 g of bodyweight) immediately prior to baseline, and approximately 12-15 minutes prior to photothrombosis. Solutions were freshly prepared, sonicated, then sterile filtered immediately prior to use. The light source for photothrombosis (561 nm, LRS-0561-GFO-00100-03, LaserGlow Technologies) was connected to the implanted ferrule (400 μm core diameter, MFC_400/430-0.48_2.5 mm_MF2.5_FLT, Doric Lenses). Brain volumes immediately below the fiberoptic cannula were illuminated for 1.5 or 8 min with 4, 5, or 6 mW intensity, where indicated.

Neuronal GCaMP6f in ipsilesional and contralesional CA1 from freely behaving Thy1-GCAMP6f mice was recorded using fiber photometry for 40 minutes surrounding the induction of photothrombosis (baseline: 10 minutes prior to 561 nm illumination, post-photothrombosis: 30 minutes following onset of illumination). Mice were excluded from the study if no GCaMP6f events were observed during baseline (GCaMP6f-negative, n = 3). For comparison between awake and anesthetized conditions, anesthetized mice were maintained under isoflurane anesthesia for the duration of the experiment (induction: 5%, maintenance: 1%).

To record neuronal GCaMP6f, a Doric fiber photometry system was used, consisting of two excitation LEDs (405 nm and 465 nm, 30 μW) controlled by the LED driver and console running DoricStudio software (Doric). Both modulated LEDs (lock-in amplification) and the 561 nm laser were filtered through a Doric Mini Cube filter set (FMC6_IE(400-410)_E1(460-490)_F1(500-540)_E2(555-570)_F2(580-880)_S) with the excitation light directed to the implanted cannula using a mono fiberoptic patchcord (Doric MFP_400/460/900-0.48_2m_FC/MF2.5). Emitted fluorescence signal between 500-540 nm was then demodulated using lock-in amplification and detected by a photoreceiver (NewPort model 2151). Following stroke, mice were followed for behavioral recovery or sacrificed for lesion histology at 48 h or 7 days post-stroke.

### Fiber photometry data analysis

Frequency-modulated GCaMP6f fluorescence (405-/465-nm excitation) was acquired at a sampling rate of 1 kHz and resampled to 100 Hz in the DoricStudio software. These data were then exported to MATLAB R2023a (Mathworks) for analyses using a custom-written script. Here, an exponential decay was fit to and then removed from both the 405 and 465 signals to account for any photobleaching-related artifacts. Motion artifacts in the photometry trace were removed by subtracting the Ca^2+^-independent 405-nm channel from the Ca^2+^-dependent 465-nm channel. The output of these transformations generated a motion-corrected dataset for change in GCaMP6f fluorescence. High frequency components of the trace were removed by smoothing the signal to 10 Hz. We normalized the signal between sessions using a modified z-score: ΔF/F = (F-F_0_)/ σF, where F is the signal, F_0_ is the median signal of baseline, and σF is the median absolute deviation of the signal. For peak GCaMP6f analyses, a low-pass filter (0.1 Hz) was used, and dysregulation was determined when ΔF/F > 20. Peak GCaMP6f was the maximum intensity value across the 2400 s recording.

To estimate the peak derivative of the GCaMP6f signal for mouse *i*, *x^i^*(*t*), the signal was first smoothed with a lowess smoothing filter and with a 10 second window, as implemented by the MATLAB (2023a) function *smooth*. This creates a smooth estimate, *x^i^*(*t*) of the signal *x^i^*(*t*) which filters much of the high-frequency jitter common in experimentally recorded signals. The derivative was then estimated with the first order finite-difference approximation:

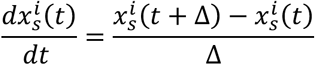

Where Δ = 0.05 seconds is the sampling frequency of the recordings. The approximation to the derivative was used to determine when the GCaMP6f signal had its maximal rise time, defined by:

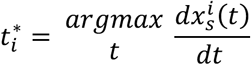

To avoid boundary effects, the derivatives 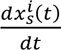 were manually set to zero in the first and last 5 seconds of recordings.

### LFP recording and analysis

Either contralesional hippocampal activity alone or bilateral hippocampal activity was recorded at 512 Hz using an AC amplifier (A-M systems) and collected using DATAQ acquisition software simultaneously during bilateral GCaMP6f recording (Compumedics Neuroscan). In experiments to blunt SD, MK-801 (0.5 mg/kg) or equivalent volume of vehicle (PBS) were added to 1% Rose Bengal solution prior to injection i.p. Bilateral GCaMP6f photometry and contralesional LFP were recorded as in previous experiments with the mouse behaving in an open field within a Faraday cage to eliminate extrinsic noise. A notch filter was used to remove artificial noise that was consistent across the recording at a specific frequency (ie. 60 Hz). Electrographic seizures were identified (> 1 s) using a custom-written MATLAB analysis script. Mean signal intensity of a 60 s noise-free segment of the baseline LFP was determined, a threshold for putative seizure activity was established as peaks > 4 standard deviations of the mean, and segments of the LFP that met this criterion were automatically identified. Seizures were then manually identified based on increased power and rhythmicity of the signal. Time-varying spectrograms were computed for contralesional LFP recording using Thomson’s multi-taper method (window length 10 s, window step 1 s, bandwidth 20 Hz, number of tapers 0; Chronux toolbox, http://chronux.org; version 2.12v02, 2018). Power spectrum density (PSD) of the LFP signal was computed using a fast Fourier transform (FFT, cosine-bell window, size 256, window overlay 93.75%) on 20 or 60 s bins of continuous noise-free segments. PSD were then averaged for ∂ (0-4 Hz), ε (4-8 Hz), α (8-12 Hz), ß (12-30 Hz), and γ (30-80 Hz). Where applicable, the mean power of time-binned segments (short-time Fourier transform, STFT, 0-80 Hz, 1 s bins) were plotted, normalized to the maximum value of the contralesional signal in the specified 60 s segment.

### Peri-stroke open field behavioral state analysis

Mice were handled (> 5 min per mouse) for the four days preceding stroke, including a 1 h familiarization period in the experimental room prior to the onset of the experiment. Position of the mouse time-locked to photometry was captured in an open field (0.5 x 0.5 m) using an overhead camera recording at 20 fps. Video analysis of behavioral state was conducted blind to the experimental conditions using a macro for Microsoft Excel (see ^150^). Sitting was defined when the mouse made no movement around its environment. Ambulating was identified when the mouse changed its location or when its hands shifted during a turn in position. Rearing was defined when the mouse stood up on its hindlegs and contacted the wall of the open field. Grooming occurred when the mouse used its hands to clean itself. Normalized photometry signal (ΔF/F; downsampled to 20 Hz) was then aligned to behavioral state data over the duration of the recording (2400 s) and exported to MATLAB for analyses using a custom-written script. For stroke mice, the average time spent in each behavioral state was determined for the intervals in before (-300, 0 s) and after (0, 300 s) the peak-derivative of ipsilesional GCaMP6f (*pre-TSD, post-TSD,* respectively). Behavioral states over time were displayed in the form of a horizontal Gantt plot, where GCaMP6f signal and behavioral state Gantt plots were shifted in time so that mice within a group were aligned to either ipsilesional TSD or centered on the CL SD peak. In shams, GCaMP6f did not display a large change in amplitude, as such, behavioral states were compared in two 300 second intervals before/after the average time of ipsilesional peak derivative in the stroke group (TSD onset, 800 s). For behavioral state linked to contralesional SD, a 100 s epoch encompassing the contralesional SD (where ΔF/F > 20) was used as the *during* epoch. The 300 s period before and after the *during* epoch were used as the *pre-CL SD* and *post-CL SD* epochs, respectively. For shams, the average time of the CL SD peak in stroke mice (1350-1450 s) was used to center the *pre-CL, during,* and *post-CL* epochs. Ambulating and rearing were combined to determine time spent active (open field activity).

### Positional analyses

Position data were obtained by annotating the center of mass of the mouse in 20-40 frames per training video using DeepLabCut^71^ to then predict the position for all video frames for all recordings, then low-pass filtered below 10 Hz using a 9-order Chebyshev type 2 filter to remove high frequency noise. Distance travelled was determined for sham and stroke mice in *pre-TSD* and *post-TSD* epochs, as defined in behavioral state analysis, and the difference in total distance was subtracted to quantify relative locomotion. Contralesional analysis used epochs of different absolute times, therefore average speed was used as a metric of locomotion. Here, distance travelled over *pre-CL, during,* and *post-CL* epochs was divided by time (s) of each epoch; difference in speed was then compared to the *pre-CL* epoch.

### Contextual fear conditioning

Contextual fear conditioning was evaluated using ANY-maze Video Tracking software. For *peri-stroke contextual fear conditioning*, on Day 1, 10 minutes prior to induction of stroke or sham, mice were placed into context 1 (transparent walls, metal grate floor, 80 lumens) where they received 3 foot shocks evenly spaced over 7 minutes (t = 140 s, t = 280 s, t = 520 s). Mice were returned to context 1 at 24 h post-stroke for 7 minutes absent of foot shocks. Percentage of time freezing (Freezing score < 50 for > 250 ms) was compared for sham and stroke (± SD) between the epoch prior to the initial foot shock on Day 1 (pre-shock) and Day 2 (*24 h post-stroke*). At 48 h post-stroke, mice were placed into a novel context (black striped walls, white solid floor) for 7 minutes absent of foot shock to evaluate generalization. For *post-stroke contextual fear conditioning*, at 6 days post-stroke, mice were placed in Context 1 and three evenly spaced foot shocks were administered over a 7-minute interval. At 7 days post-stroke, mice were returned to Context 1 absent of foot shocks. Percentage of time freezing was compared between sham and stroke (± SD) between Day 1 pre-shock relative to throughout Day 2 (7 d *post-stroke*). Data were presented as percentage of time spent freezing during the specified epoch.

### Lesion histology

In a subset of mice, at 48 h post sham illumination or photothrombosis, brain tissue viability was assessed using triphenyltetrazolium chloride (TTC). Briefly, mice were anesthetized using inhaled isoflurane, decapitated, and brains extracted into ice-cold saline for 5 min. Coronal slices (2 mm) were made using a BrainBlock (Zivic Instruments), then placed in a 1% TTC solution for 30 min at 37 °C. Slices were then moved to 4% paraformaldehyde (PFA) for 24 h prior to imaging. Digital images were acquired with a desktop color scanner (600 dpi) and lesion area (white) relative to healthy tissue (red) was analyzed using FIJI. At 7 days post unilateral photothrombosis, brains were collected to determine the histological profile of the lesion for each experimental group. Mice were injected with sodium pentobarbital (30 mg/kg, i.p.) then transcardially perfused with saline and 4 % PFA in succession. Extracted brains were placed in 4 % PFA for > 24 h, then in 30 % sucrose in PBS at 4 °C. After fixation, 40 µm thick brain slices were collected in the region surrounding the fiberoptic implant for both control and stroke experimental groups using a cryostat and mounted on slides (Fisher Scientific). Slices were rehydrated in PBS, permeabilized with 0.3 % Triton-X (PBS-Tx) for 10 min, then blocked in 10 % normal goat serum (NGS) in PBS-Tx for 1 h. Slices were incubated in anti-Iba1 (1:500, Invitrogen #PA5-27436), Cy3-conjugated mouse anti-GFAP (1:500, Sigma #C9205) for 24 h at room temperature. Slices were then washed in PBS-Tx and incubated in AlexaFluor647 goat anti-mouse (1:500, Invitrogen #A32733) for 2 h at room temperature. Both primary and secondary antibody incubations were done in PBS-Tx (+ 2% NGS). Slices were then washed in PBS-Tx, then PBS, prior to mounting coverslips with VectaShield + DAPI (Vector Labs #H-1200-10). Images were captured on an Olympus VS110 Slide Scanner. Area of Iba1 and GFAP reactivity were evaluated by thresholding to where hippocampal intensity of each signal was zero in sham mice, the area of positive pixels in the ipsilesional hemisphere was then determined. For GCaMP6f expression, the hippocampus was outlined based on anatomical structure in the DAPI channel, the percentage of pixels that were GCaMP6f-positive were then determined.

### Chronic window surgery with fiberoptic implant

Chronic window surgeries were adapted from previously described methods^86,87^, performed here in female Thy1-GCaMP6f mice (n = 2 without hippocampal SD, n = 3 with hippocampal SD, n = 1 with cortical SD). Mice were isoflurane anesthetized (4 % induction, 1.5-2.5 % maintenance, 0.5 L/min oxygen) and Meloxicam (5 mg/kg) was administered subcutaneously for analgesia. Body temperature was maintained at 37 °C, eyes were lubricated (Opticare, CLC Medica, Canada), and bupivacaine (intradermal, 0.05 ml, 5 mg/ml) was administered locally at the excision site. Following disinfection with 3 x alternating chlorhexidine (2 %) and alcohol (70 %), the skull was exposed with a skin excision from 3 mm anterior to bregma to 2 mm posterior to lambda, and bilaterally to the temporalis muscles. Fiberoptic implants (Doric Lenses, identical to above) aimed at the dorsal hippocampus were implanted bilaterally, angled in two planes to facilitate simultaneous widefield cortical imaging. Bore holes were drilled at AP -3.0 mm, ML +/- 2.0 mm, and ferrules inserted 30° off the vertical plane and 20° off the sagittal plane, extending 1.4 mm longitudinal to a final target of AP -2.3 mm, ML +/- 1.75 mm, DV -1.2 mm relative to the cortical surface. Ferrules were fixed in place with dental cement (C&B-Metabond, Parkell). A flat 6 x 8 mm glass coverslip (tapered at the anterior end by 2 mm) was subsequently fixed to the skull with transparent dental cement. A metal screw was fixed to the posterior edge of the implant between ferrules to facilitate head fixation. Mice recovered for 7 days prior to further interventions, allowing for full cement hardening and wound healing.

### Mesoscale imaging with bilateral fiber photometry

Mice were habituated to handling and head fixation with the embedded screw over 5 days. Images were acquired using a Quantalux 2.1 MP Monochrome sCMOS Camera (Thorlabs) at 20 Hz (49.7 ms exposure time) with a 256 x 256-pixel resolution (26.2 px/mm) through a Nikon 55 mm lens (f/2.8 aperture) with a 535/30 emission filter. GCaMP6f was excited with a 475 nm LED (472/30 filter) attached to an articulating arm (10-15 mW/cm^2^). Focus was set ∼500 µm below the cortical surface to minimize signal distortion from large blood vessels. Illumination and frame capture was controlled using commercial software (Labeo Technologies, Inc). To facilitate Rose Bengal-mediated photothrombosis, reflectance imaging that would enable hemodynamic correction was not collected. A TTL trigger input to the image acquisition control unit signalling the start and end of photometry signal acquisition enabled precise alignment of photometry and widefield cortical imaging.

### Mesoscale image analysis

Image time-stacks were analyzed using custom-written MATLAB code (Mathworks, MA). First, stacks were truncated to temporally align to the photometry signal, and pixel responses were expressed as a change in fluorescence (ΔF/F_0_) relative to the 5 min baseline of the recording. As fiberoptic implants partially occluded the posterior cortex, 26 ROIs were selected (13 in each hemisphere) based on manual alignment to the Allen Brain Atlas. 11 x 11 pixel regions were averaged for each ROI to estimate cortical Ca^2+^ fluctuations from each region. To facilitate analysis of SD propagation following photothrombosis invading superficial cortical tissue, 5 ROIs were selected at even intervals (1.2 mm) from ischemic core along the propagation direction axis of the SD.

### Analysis & Statistics

GraphPad Prism 9.0 software was used for statistical analyses. Statistical tests for each experiment are described in the figure legends (data presented as mean ± s.e.m.). For pairwise comparisons, two-sided unpaired t-tests, paired t-tests, or Mann-Whitney tests were performed based on normality of the data. For data where more than two groups were compared, either one-way ANOVA or two-way ANOVAs were performed with Bonferroni or Tukey *post-hoc* tests, where appropriate. In the case of repeated measures, a repeated measures two-way ANOVA was performed. The number of biological replicates and p-values are described in the figure legends.

### Data availability

Data available upon reasonable request. Source data are included with this paper.

### Code availability

The scripts (v0.0.2) used to process and analyse fiber photometry and behavioral data are available at https://doi.org/10.5281/zenodo.8303055.

## Acknowledgments

We thank Cheryl Breiteneder and Rodney Barasi for technical assistance and Núria Daviu Abant and Adam Institoris for careful review of the manuscript. We thank the Cumming School of Medicine Optogenetics Platform for fiber photometry training. We thank SciDraw for use of open-source drawings. This research was funded by operating grants to R.J.T. from the Canadian Institute for Health Research (PJT 169174 & PJT 168968) and the Krembil Foundation and to G.C.T. from a Natural Sciences & Engineering Research Council of Canada (NSERC) Discovery Grant (RGPIN/04140-2019). AKJB was supported by an Alberta Innovates (2+1) postdoctoral fellowship and a Donald Burns & Louise Berlin Postdoctoral Fellowship in Dementia Research.

## Author Contributions

AKJB & RJT designed the study. AKJB performed all *in vivo* stroke & behavior recordings, wrote LFP analysis scripts, analyzed data, prepared figures, and wrote/edited the manuscript. DMA implanted mesoscale imaging windows and performed mesoscale and FP experiments with AKJB. YF & CMS performed immunohistochemistry. RCG performed electrode implants and assisted with LFP recording. LM wrote photometry and DLC analysis scripts. TF created behavior state analysis macro. CE analyzed some behavioral data. WN wrote behavioral state analysis script. RJT, AM, & GCT supervised the study and edited the manuscript.

