## Supplementary Information for "Contralesional hippocampal spreading depolarization promotes functional recovery after stroke"

**Supplement: Detailed Statistical Information**

Figure 1.

| Figure Panel | Response variable | n | statistical analysis | Result |
|---|---|---|---|---|
| Figure 1e | Peak GCaMP6f (ΔF/F) | sham - female = 7, stroke - female = 32 , sham - male = 6, stroke male = 20 | two-way ANOVA with Tukey *post hoc* | sex: F1, 61 = 0.20, p = 0.65; sham vs. stroke: F1, 61 = 43.74, **** p < 0.0001; interaction: F1,61 = 0.20, p = 0.65; female – stroke: 1205 arb. units ± 106.8, n = 32; male – stroke: 1051 arb. units ± 111.7 |
| Figure 1f | Peak derivative (GCaMP6f) | stroke - female = 14, stroke - male = 10 | unpaired t-test | p = 0.6887, female: 0.29 ± 0.04; male: 0.32 ± 0.07 |
| Figure 1g | Time to TD | stroke - female = 31, male = 18 | unpaired t-test | p = 0.9355 (female: 221.5 s ± 18.3; male: 219.2 s ± 18.3) |
| Figure 1h | Lesion size (mm2) | sham - female = 7, stroke - female = 13 , sham - male = 6, stroke male = 10 | Two-way ANOVA with Tukey *post hoc* | sex: F1, 32 = 0.47, p = 0.50; sham vs. stroke: F1, 32 = 31.65, **** p < 0.0001; interaction: F1,32 = 0.51, p = 0.48 |
| Figure 1i | Peak GCaMP6f (ΔF/F)-Lesion Size (mm2) | sham - female = 7, stroke - female = 13 , sham - male = 6, stroke male = 10 | Simple linear regressions | female: $R^2$ = 0.36, ** p = 0.006; male: $R^2$ = 0.54, ** p = 0.003 |
| Figure 1m | Open field activity (Δ) | sham - females = 7, stroke - females = 7, sham - males = 5, stroke - males = 9 | Two-way ANOVA with Tukey *post hoc* | sex: F1, 26 = 0.15, p = 0.70; sham vs. stroke: F1, 26 = 22.32, **** p < 0.0001; interaction: F1, 26 = 0.11, p = 0.74 |
| Figure 1o | Distance travelled (Δ) | sham - females = 6, stroke - females = 8 | unpaired t-test | * p = 0.0186 (sham: -0.56 m ± 0.66; stroke: +5.01 m ± 1.69) |
| Figure 1q | Distance travelled (Δ) | sham - males = 5, stroke - males = 10 | unpaired t-test | ** p =0.0039 (sham: -2.26 m ± 0.84; stroke: +3.70 m ± 1.11) |

Figure 2.

| Figure Panel | Response variable | n | statistical analysis | Result |
|---|---|---|---|---|
| Figure 2d | Peak GCaMP6f (CL SD) | n = 20 females, n = 5 males | Mann-Whitney | p = 0.097, female: 742.6 arb. units ± 84.3, male: 506.0 arb. units ± 135.2, |
| Figure 2g | CL events (per min) | n = 25 | paired t-test | **** p < 0.0001., pre-CL SD: 7.3 ± 0.84, post CL-SD: 2.56 ± 0.66, |
| Figure 2h | Peak GCaMP6f (CL SD) - CL events/min | n = 25 | simple linear regression | female: $R^2$ = 0.24, * p = 0.01 |
| Figure 2l | Open field activity (females) | sham, n = 6, stroke, n = 10 | unpaired t-tests | ** p < 0.01, * p < 0.05, sham – during: -16.6% ± 4.75, stroke – during: +22.89% ± 11.0, sham – post: -12.14% ± 4.5, stroke – post: +20.4% ± 5.61, |
| Figure 2m | Open field activity (males) | sham n = 5, stroke n = 2 | unpaired t-tests | n.s.; sham – during: -9.31% ± 5.4, stroke – during: +9.1% ± 5.1, sham – post: -6.94% ± 6.43, stroke – post: +4.26% ± 10.34 |
| Figure 2o | Speed (females) | sham, n = 6, stroke, n = 10 | unpaired t-tests | p = 0.0275, sham – during: -0.002 m/s ± 0.007; stroke – during: 0.006 m/s ± 0.006; sham – post: -0.002 m/s ± 0.0.005; stroke – post: 0.01 m/s ± 0.003 |
| Figure 2p | Speed (males) | sham n = 5, stroke n = 2 | unpaired t-tests | sham – during: -0.01 m/s ± 0.003, stroke – during: -0.002 m/s ± 0.005, sham – post: -0.006 m/s ± 0.0.003, stroke – post: 0.001 m/s ± 0.003 |
| Figure 2q | Distance (post-TSD) | stroke - SD, n = 10; stroke + SD, n = 8 | unpaired t-test | ** p < 0.01, -SD: 12.12 ± 1.87 m in 20 min; +SD: 28.71 ± 5.99 m |

Figure 3.

| Figure Panel | Response variable | n | statistical analysis | Result |
|---|---|---|---|---|
| Figure 3b | Freezing time (%) | sham, n = 4, stroke - SD, n = 3, stroke + SD, n = 3 | Two-way RM ANOVA with Bonferroni post hoc | time: $F_{1,19}$ = 69.59, **** p < 0.0001; insult: $F_{2, 19}$ = 37.41, **** p < 0.0001; interaction: $F_{2, 19}$ = 34.68, **** p < 0.0001]) |
| Figure 3c | Freezing time (%) | sham, n = 6, stroke - SD, n = 7, stroke + SD, n = 8 | Two-way RM ANOVA with Bonferroni post hoc | time: F(1,7) = 18.44, ** p < 0.01; insult: $F_{2, 7}$ = 4.8, * p < 0.05; interaction: $F_{2, 7}$ = 4.92, * p < 0.05] |
| Figure 3d | Peri-stroke: Peak GCaMP6f (IL TSD)-Freezing (Day 2) | females, n = 15 | simple linear regression | $R^2$ = 0.30, p = 0.033 |
| Figure 3e | Peri-stroke: Peak GCaMP6f (CL SD)-Freezing (Day 2) | females, n = 15 | simple linear regression | $R^2$ = 0.034, p = 0.51 |
| Figure 3g | Freezing time (%) | sham, n = 4 | paired t-test | p = 0.0233 |
| Figure 3h | Freezing time (%) | stroke - SD, n = 3 , stroke + SD, n = 3 | Two-way RM ANOVA with Bonferroni post hoc | time: $F_{1,11}$ = 60.82, **** p < 0.0001; insult: $F_{1, 11}$ = 23.94, *** p = 0.0005; interaction: $F_{1, 11}$ = 25.27, *** p = 0.0004. |
| Figure 3i | Freezing time (%) | stroke - SD, n = 6 , stroke + SD, n = 7 | Two-way RM ANOVA with Bonferroni post hoc | time: F1,4 = 16.5, p = 0.02; insult: F1, 4 = 0.005, p = 0.95; interaction: F1, 4 = 2.116, p = 0.2195 |
| Figure 3j | Post-stroke: Peak GCaMP6f (IL TSD)-Freezing (Day 2) | females, n = 14 | simple linear regression | $R^2$ = 0.0005, p = 0.94 |
| Figure 3k | Post stroke: Peak GCaMP6f (CL SD)-Freezing (Day 2) | females, n = 14 | simple linear regression | $R^2$ = 0.42, p = 0.011 |

Figure 4.

| Figure Panel | Response variable | n | statistical analysis | Result |
|---|---|---|---|---|
| Figure 4e | Power (dB) | n = 5 females | RM two-way ANOVA with Bonferroni post-hoc | (frequency: $F_{3.62, 58.0}$ = 189.6, **** p < 0.0001; epoch: $F_{3, 16}$= 23.14, **** p < 0.0001; interaction: $F_{366, 1952}$ = 6.95, **** p < 0.0001) |
| Figure 4f | Power (dB) | n = 5 females | RM two-way ANOVA with Bonferroni *post-hoc* | (frequency: F2.61, 57.36 = 80.25, **** p < 0.0001; epoch: F2, 22 = 1.88, p = 0.18; interaction: F256, 2816 = 2.53, **** p < 0.0001) |
| Figure 4i | Power (dB) | n = 4 males | RM two-way ANOVA with Bonferroni *post-hoc* | (frequency: F128,1536 = 80.25, **** p < 0.0001; epoch: F1, 12 = 3.61, p = 0.082; interaction: F128, 1536 = 2.65, **** p < 0.0001) |
| Figure 4k | Time (s, FP) vs. Time (s, LFP) | n = 5 CL SDs from 4 female mice | Linear regression | onset: R2 = 0.99, **** p < 0.0001; end: R2 = 0.99, **** p < 0.0001 |
| Figure 4l | Cumulative seizure duration | female, n = 5; male, n = 4 | unpaired t-test | p = 0.32, female: 78.32 s ± 23.9, n = 5; male: 44.4 s ± 19.31 |

Figure 5.

| Figure Panel | Response variable | n | statistical analysis | Result |
|---|---|---|---|---|
| Figure 5d | Power (normalized) | n = 4 females | two-way RM ANOVA with Bonferroni *post hoc* | time: $F_{119, 357} = 1.72$, **** $p < 0.0001$; hemisphere: $F_{1, 3} = 347.2$, *** $p = 0.0003$; interaction: $F_{119, 357} = 1.59$, *** $p = 0.0006$, |
| Figure 5e | Power (dB) - ipsilesional | n = 4 females | two-way RM ANOVA with Bonferroni *post hoc* | time: F (119, 357) = 44.37, p < 0.0001; epoch: F (1, 3) = 31.26, p = 0.0113; Interaction: F (119, 357) = 6.783, p < 0.0001 |
| Figure 5f | Power (dB) - contralesional | n = 4 females | two-way RM ANOVA with Bonferroni *post hoc* | time: F (119, 357) = 13.77, P<0.0001; epoch: F (1, 3) = 0.8182, P=0.4324; interaction: F (119, 357) = 1.207, P=0.0966 |

Figure 6.

| Figure Panel | Response variable | n | statistical analysis | Result |
|---|---|---|---|---|
| Figure 6i | Peak GCaMP6f (IL TSD) | vehicle, n = 7; MK801, n = 6 | unpaired t-test | vehicle: 1360 arb. units ± 174.7; MK-801: 879.6 arb. units ± 97.11; * p = 0.0429 |
| Figure 6k | Peak GCaMP6f (CL SD) | vehicle, n = 6; MK801, n = 6 | unpaired t-test | vehicle: 724.1 arb. units ± 76.5; MK-801: 246.1 arb. units ± 20.84; **** p < 0.0001 |
| Figure 6l | Seizure amplitude | vehicle - SD, n = 4; vehicle + SD, n = 5, MK801 n = 6 | one-way ANOVA with Bonferroni post hoc | vehicle - SD: 0.054 µV ± 0.016; vehicle + SD: 0.044 µV ± 0.007; MK-801: 0.15 µV ± 0.041; $F_{2,12} = 4.49$, * p = 0.035 |
| Figure 6m - pre | Power (dB) | vehicle, n = 5; MK801, n = 9 | RM two way ANOVA with Bonferroni post hoc | frequency: $F_{4.44, 53.3} = 74.7$, **** p < 0.0001; treatment: $F_{1, 12} = 10.91$, ** p = 0.0063; interaction: $F_{128, 1536} = 6.70$, **** p < 0.0001 |
| Figure 6m - seizure | Power (dB) | vehicle, n = 5; MK801, n = 10 | RM two way ANOVA with Bonferroni post hoc | frequency: $F_{4.35, 52.2} = 105.7$, **** p < 0.0001; treatment: $F_{1, 12} = 14.77$, ** p = 0.0023; interaction: $F_{128, 1536} = 4.42$, **** p < 0.0001 |
| Figure 6m - SD | Power (dB) | vehicle, n = 5; MK801, n = 11 | RM two way ANOVA with Bonferroni post hoc | frequency: $F_{4.64, 55.7} = 39.74$, **** p < 0.0001; treatment: $F_{1, 12} = 10.91$, ** p = 0.0063; interaction: $F_{128, 1536} = 4.46$, **** p < 0.0001 |
| Figure 6m - post | Power (dB) | vehicle, n = 5; MK801, n = 12 | RM two way ANOVA with Bonferroni post hoc | frequency: $F_{3.60, 43.14} = 63.25$, **** p < 0.0001; treatment: $F_{1, 12} = 17.96$, ** p = 0.0012; interaction: $F_{128, 1536} = 8.24$, **** p < 0.0001 |
| Figure 6o | Cumulative seizure duration | vehicle - SD, n = 4; vehicle + SD, n = 5, MK801, n = 6 | one-way ANOVA with Bonferroni post-hoc | vehicle - SD: 40.6 s ± 19.6, n = 4; vehicle + SD: 87.5 s ± 6.93, n = 5; MK-801: 295.7 s ± 73.7, n = 6; , $F_{2, 12} = 6.895$, * p = 0.01 |

| | | | | |
|---|---|---|---|---|
| Figure 6q | Freezing % | vehicle - SD, n = 4; vehicle + SD, n = 7; MK801 - sham, n = 4; MK801 - stroke, n =7 | Two way ANOVA with Bonferroni post hoc | time: $F_{1, 18}$ = 20.28, *** p = 0.0003; treatment: $F_{3, 18}$ = 2.66, p = 0.08; interaction: $F_{3, 18}$ = 2.98, p = 0.06 |
