## Extended Data Figures for "Contralesional hippocampal spreading depolarization promotes functional recovery after stroke"

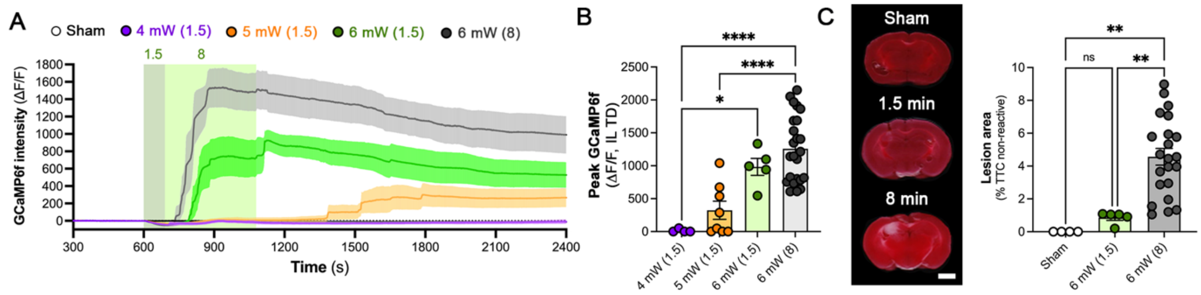


**Extended Data Figure 1. Illumination power and time-dependent increases in ipsilesional TD and lesion size. (A)** Increasing laser illumination power and duration increases neuronal $Ca^{2+}$ influx (GCaMP6f, ΔF/F) during unilateral hippocampal photothrombosis in awake mice (mean ± s.e.m.). Distinct laser illumination parameters were used to initiate photothrombosis through the optical fiber also utilized for photometry (561 nm – black: 6 mW, 8 min, n = 24; green: 6 mW, 1.5 min, n = 5; orange: 5 mW, 1.5 min, n = 8; purple: 4 mW, 1.5 min, n = 4; grey: sham, n = 4). **(B)** Increasing illumination duration and power increased ipsilesional TD (peak GCaMP6f, ΔF/F; 6 mW, 8 min, 1258.0 ± 102.8; green: 6 mW, 1.5 min, 982.6 a.u. ± 128.5; orange: 5 mW, 1.5 min, 323.4 a.u. ± 138.4; purple: 4 mW, 1.5 min, 13.0 a.u. ± 13.0, one-way ANOVA with Tukey *post hoc*, *p < 0.05, **p < 0.01, ****p < 0.0001). **(C)** Photothrombotic stroke lesion scales with illumination duration (representative TTC-stained coronal sections). Scale bars, 2 mm. Stroke lesions (% TTC-negative brain area) are larger in 8 min illuminated mice. One-way ANOVA with Tukey *post hoc*, sham: 0.0% ± 0.0, stroke (1.5 min, 6 mW): 0.86% ± 0.17, stroke (8 min, 6 mW): 4.57% ± 0.17 at 48 h post-stroke; ****p < 0.0001.

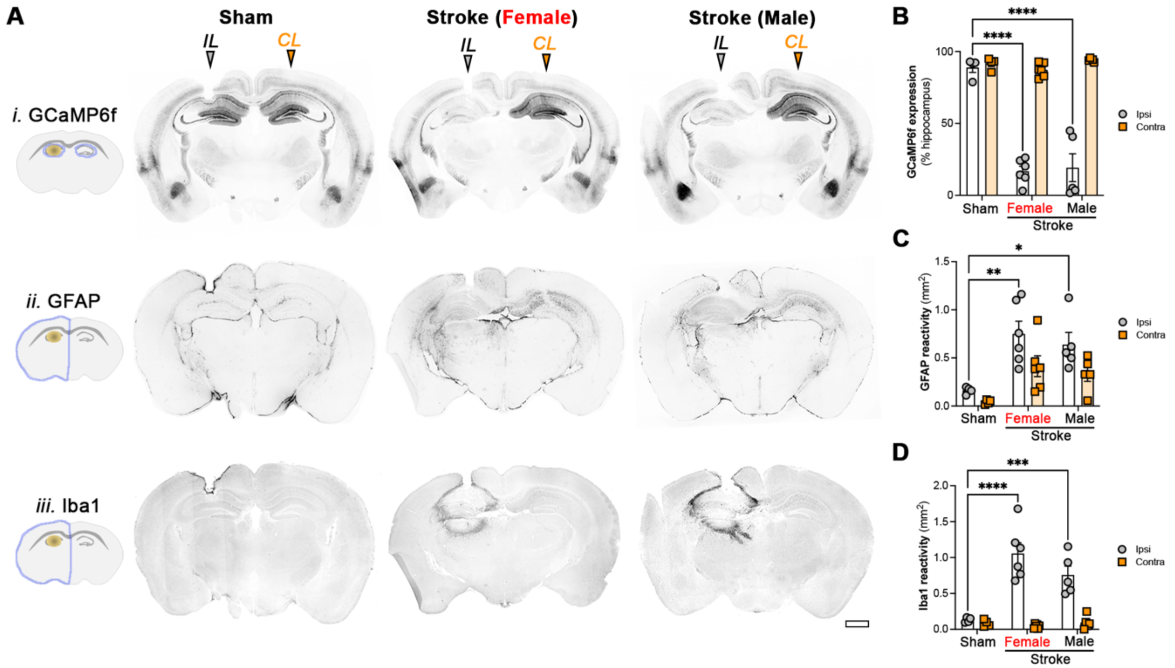


**Extended Data Figure 2. Neuronal loss and glial reactivity are indistinguishable between males and females after unilateral hippocampal stroke. (A)** Representative coronal sections highlight reduction of (**i**) hippocampal GCaMP6f expression and increased (**ii**) GFAP and (**iii**) Iba1-immunoreactivity in the ipsilesional hemisphere of female (n = 6) and male (n = 5) mice below the optical fiber tract one week following unilateral hippocampal photothrombosis relative to mice experiencing sham illumination (n = 4). Scale bars, 1 mm. **(D)** GCaMP6f expression (% of hippocampus positive for GCaMP6f) is reduced in the ipsilesional hippocampus after stroke (sham, ipsi: 88.91 ± 3.39%, sham, contra: 91.67 ± 2.07%; stroke – female, ipsi: 16.78% ± 3.5, stroke – female, contra: 87.23 ± 1.97%; stroke – male, ipsi: 19.24 ± 9.7%, stroke – male, contra: 94.10 ± 0.53%; two-way ANOVA with Bonferroni *post-hoc,* stimulus: $F_{2,24} = 36.65$, $p < 0.0001$, hemisphere: $F_{1,24} = 167.3$, $p < 0.0001$, interaction: $F_{2,24} = 33.64$, $p < 0.0001$; **** $p < 0.0001$). **(E)** GFAP reactivity in the ipsilesional hemisphere is increased following unilateral hippocampal photothrombosis (sham, ipsi: 161.3 ± 18.5 $\mu m^2$, sham, contra: 42 ± 12 $\mu m^2$; stroke – female, ipsi: 749.7 ± 131.3 $\mu m^2$, stroke – female, contra: 414 ± 108 $\mu m^2$, stroke – male, ipsi: 517.3 ± 47.3 $\mu m^2$, stroke – male, contra: 335 ± 79 $\mu m^2$; two-way ANOVA with Bonferroni *post-hoc,* stimulus: $F_{2,24} = 10.66$, $p = 0.0005$, hemisphere: $F_{1,24} = 8.52$, $p = 0.0075$, interaction: $F_{2,24} = 0.56$, $p = 0.58$; * $p < 0.05$, ** $p < 0.01$). **(F)** Iba1 reactivity ($mm^2$) in the ipsilesional hemisphere is increased following unilateral hippocampal photothrombosis (sham, ipsi: 127.3 ± 17.0 $\mu m^2$, sham, contra: 85 ± 24 $\mu m^2$; stroke – female: 1057.0 ± 150.1 $\mu m^2$, stroke – female, contra: 44 ± 12 $\mu m^2$, stroke – male: 982.6 ± 206.9 $\mu m^2$, stroke – male, contra: 96 ± 42 $\mu m^2$; two-way ANOVA with Bonferroni *post-hoc,* stimulus: $F_{2,24} = 11.13$, $p = 0.0004$, hemisphere: $F_{1,24} = 55.49$, $p < 0.0001$, interaction: $F_{2,24} = 13.03$, $p = 0.0001$; *** $p < 0.001$, **** $p < 0.0001$).

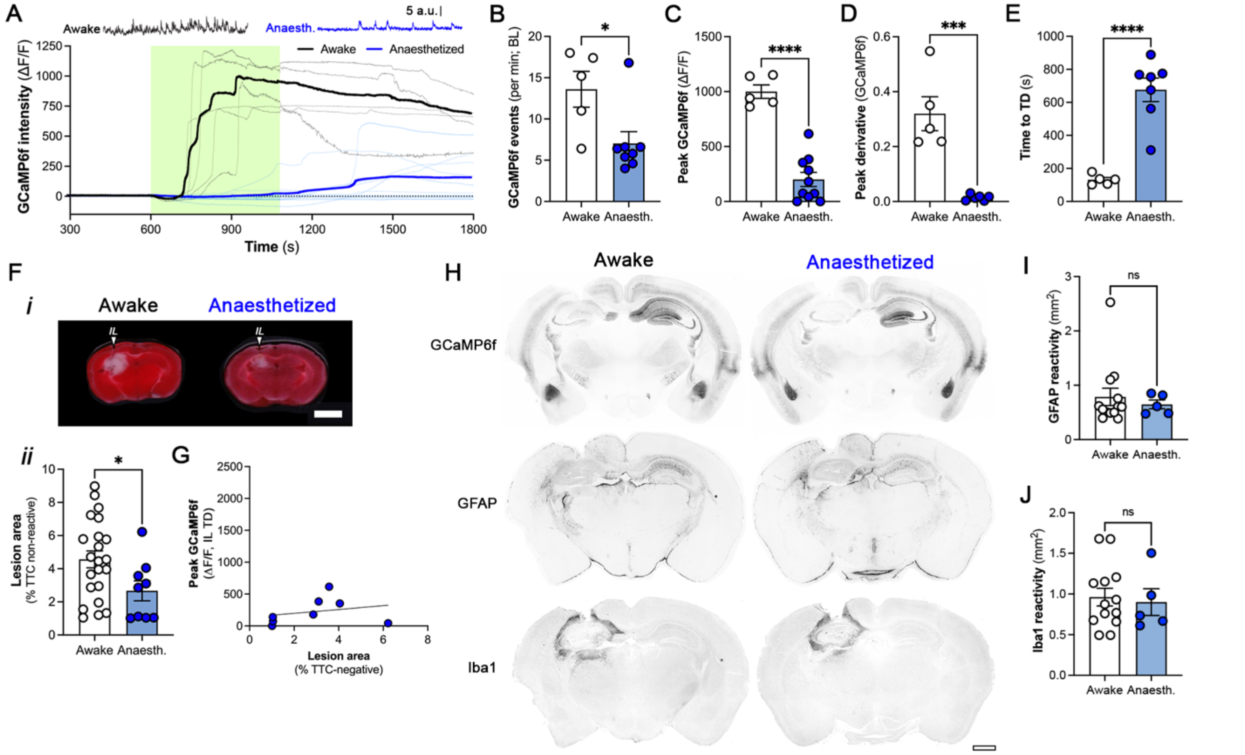


**Extended Data Figure 3. Isoflurane anesthesia protect against focal hippocampal stroke.** (A) Representative 60 s segments of baseline hippocampal GCaMP6f photometry recordings of awake (black) and anesthetized (blue) Thy1GCaMP6f mice. Neuronal $Ca^{2+}$ dysregulation ($\Delta F/F$; mean: thick lines; individual traces: thin lines) induced by unilateral hippocampal photothrombosis is distinct between awake (black) and anesthetized (blue) mice. Photothrombotic illumination (561 nm, 6 mW, 10-18 min). **(B)** Awake mice have more frequent $Ca^{2+}$ events per min than anesthetized mice during the baseline epoch (awake: 13.6 ± 2.2, n = 5; anesthetized: 7.0 ± 1.4, n = 8; unpaired t-test, *$p < 0.05$). **(C)** Awake mice have larger TD (peak GCaMP6f, $\Delta F/F$) following hippocampal photothrombosis (awake: 999.7 a.u. ± 61.0, n = 5; anesthetized: 200.5 a.u. ± 64.1, n = 8; unpaired t-test, ***$p < 0.001$). **(D)** Slope of ipsilesional TD (peak derivative of GCaMP6f) is larger in awake mice (awake: 0.32 ± 0.06, n = 5; anesthetized: 0.04 ± 0.004, n = 6; unpaired t-test, ***$p < 0.001$). **(E)** TD onset ($t_{peak\ derivative}$, GCaMP6f) is more rapid in awake mice following onset of photothrombotic illumination (awake: 131.4 s ± 13.8, n = 6; anesthetized: 676.2 s ± 71.7, n = 7; unpaired t-test, ****$p < 0.001$). **(F, *i*)** Representative TTC-stained coronal sections from awake and anesthetized mice 48 h after unilateral hippocampal photothrombosis reveal that (***ii***) awake mice have larger lesions than anesthetized mice (%TTC-negative brain area, awake: 4.57% ± 0.51, anesthetized: 2.67% ± 0.6, unpaired t-test, * $p < 0.05$; 48 h post-illumination). **(G)** No correlation exists between ipsilesional TD and hippocampal stroke lesion size of anesthetized mice ($R^2$ = 0.02, p = 0.73). **(H)** Representative coronal sections at fiber implant site reveal that glial reactivity is not significantly different between awake (n = 13) and anesthetized mice (n = 5) one-week post-stroke. Scale bars, 1 mm. **(I)** Iba1 reactivity ($mm^2$) is not different in awake and anesthetized stroke (awake: 962.0 $\mu m^2$ ± 108.8, anesthetized: 900.4 $\mu m^2$ ± 163.7, unpaired t-test). **(J)** GFAP reactivity ($mm^2$) is not different in awake and anesthetized stroke (awake: 784.7 $\mu m^2$ ± 159.8, anesthetized: 647.4 $\mu m^2$ ± 79.7, unpaired t-test).

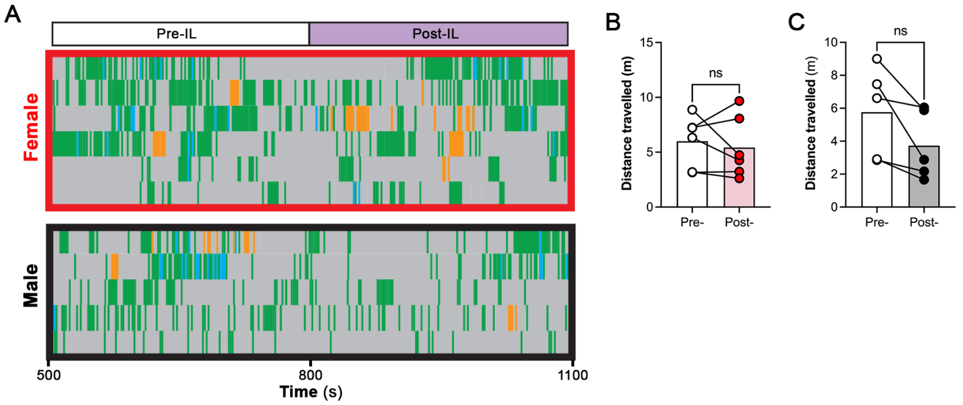


**Extended Data Figure 4. Sham mice do not alter activity or locomotion in the approximate peri-TD period. (A)** Gantt behavioral plots for all analyzed female (red outline, n = 6) and male (black outline, n = 5) sham mice centered on the average time of TD onset identified in the stroke group (800 s). **(B)** Distance travelled by female sham mice does not change following the average time of TD onset in stroke mice (800 s, n = 6; pre-IL: 6.0 m ± 0.95, post-IL: 5.43 m ± 1.15). **(C)** Distance travelled by male sham mice does not change following the average time of TD onset in stroke mice (800 s, n = 5, pre-IL: 5.77 m ± 1.24, post-IL: 3.73 m ± 0.93).

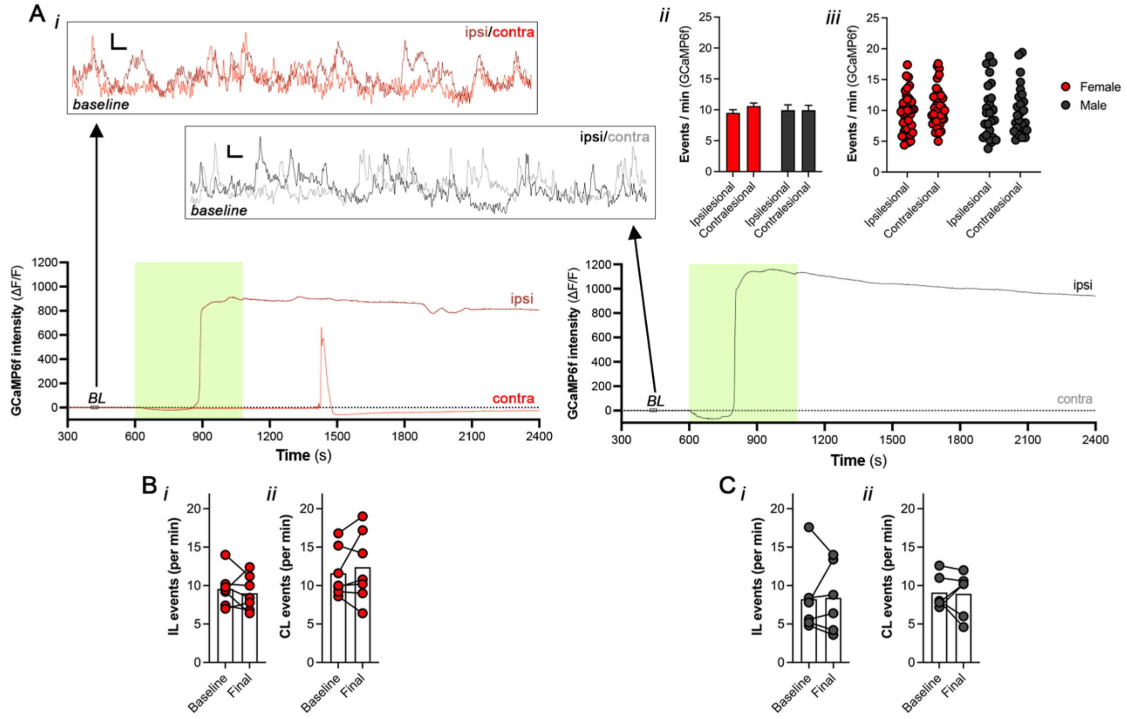


**Extended Data Figure 5. Basal ipsi- and contralesional $Ca^{2+}$ transients are unchanged over recording period. (A)** Baseline ipsilesional (dark) and contralesional (light) GCaMP6f photometry traces (scale bar: 1 arb. units, 1 s) from awake and freely-behaving **(i)** female (red) and **(ii)** male (black) mice preceding photothrombosis (see Fig. 2 for SD analyses). **(iii)** No significant differences between $Ca^{2+}$ events in ipsilesional and contralesional hippocampi between females (red, IL: 9.51 ± 0.52, CL: 10.6 ± 0.49) and males (black, IL: 9.95 ± 0.85, CL: 9.94 ± 0.78), displayed as mean +/- s.e.m. (***i***, bar graph) or individual values (***ii***, scatter plot). Two-way ANOVA with Bonferroni *post hoc*, sex: $F_{1, 63} = 0.02$, $p = 0.88$; hemisphere: $F_{1, 63} = 1.02$, $p = 0.32$; interaction: $F_{1, 63} = 1.02$, $p = 0.32$. **(B)** No change in $Ca^{2+}$ events (per min) between *baseline* (300 - 600 s) and *final* (2100 - 2400 s) epochs in **(i)** ipsilesional (BL: 9.57 ± 0.87, final: 9.0 ± 0.86, n = 7) and **(ii)** contralesional GCaMP6f (BL: 11.6 ± 1.2, final: 12.4 ± 1.72, n = 7) from female shams (paired t-tests). **(C)** No change in $Ca^{2+}$ events (per min) between *baseline* and *final* epochs in **(i)** ipsilesional (BL: 8.23 ± 1.97, final: 8.4 ± 1.84, n = 6) and **(ii)** contralesional GCaMP6f (BL: 9.1 ± 0.89, final: 8.93 ± 1.2, n = 6) from male shams. Unpaired t-tests.

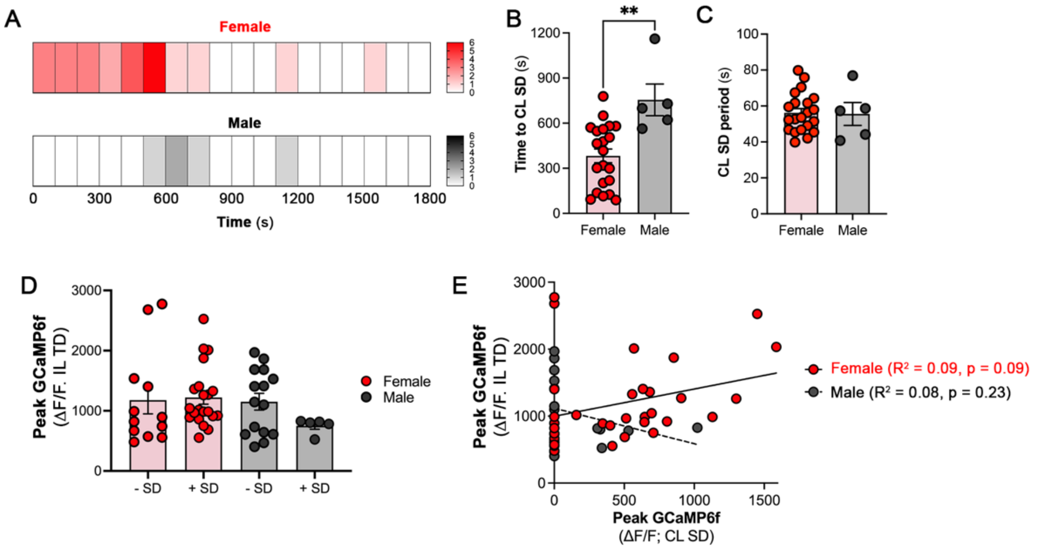


**Extended Data Figure 6. Sex-specific contralesional SD characteristics. (A)** Contralesional SD occur more rapidly in females than in males. Frequency plot (100 s bins) for CL SD onset following ipsilesional TD (female: n = 21, male: n = 5). **(B)** Onset of CL SD is more rapid in females (females: 382.6 s ± 45.71, n = 21; males: 755.0 s ± 105.6, n = 5; unpaired t-test, ** p = 0.002). **(C)** No significant difference in CL SD period (females: 56.22 ± 2.40, n = 21; males: 55.58 ± 6.40, n = 5; unpaired t-test). **(D)** Ipsilesional TD amplitude has no impact on the presence of CL SD. Two-way ANOVA with Tukey *post-hoc* (sex: $F_{1, 49} = 1.88$, p = 0.18; ± SD: $F_{1, 49} = 0.97$, p = 0.33; interaction: $F_{1, 49} = 1.50$, p = 0.23). **(E)** Ipsilesional TD and contralesional SD amplitudes (ΔF/F) are not correlated in females (red) or males (black). Linear regression (female: solid line; $R^2 = 0.09$, p = 0.09; male: dotted line; $R^2 = 0.08$, p = 0.23).

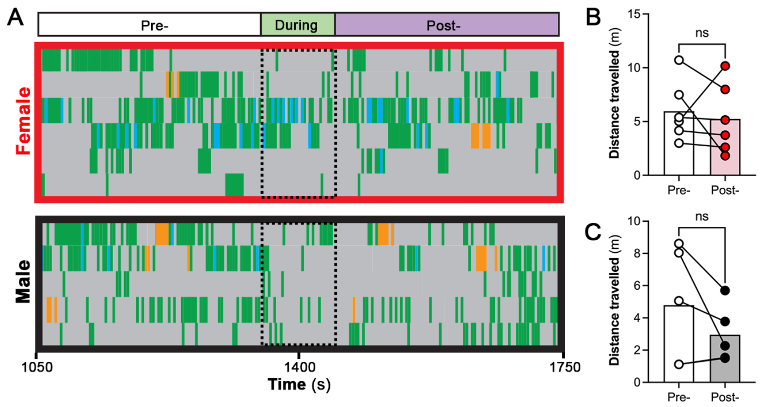


**Extended Data Figure 7. Sham mice do not alter activity or locomotion in the approximate peri-CL SD epoch. (A)** Gantt behavioral plots for all analyzed female (red outline, n = 6) and male (black outline, n = 4) sham mice centered on the average time of CL SD identified in the stroke group (1400 s). **(B)** Distance travelled by female sham mice does not change following the average time of CL SD in stroke mice (pre-: 5.96 m ± 1.13, post-: 5.24 m ± 1.33, paired t-test). **(C)** Distance travelled by male sham mice does not change following the average time of CL SD in stroke mice (pre-: 4.79 m ± 1.61, post-: 2.96 m ± 0.8, paired t-test).

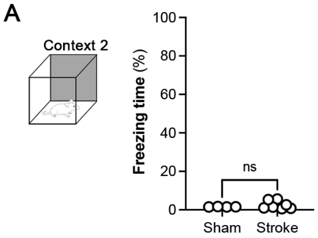


**Extended Data Figure 8. Freezing behavior is specific to fear conditioned context. (A)** No change in time spent freezing (%) by sham or stroke mice in a novel context independent of foot shock. Unpaired t-test, $p = 0.33$ (sham: n = 4, stroke: n = 8).

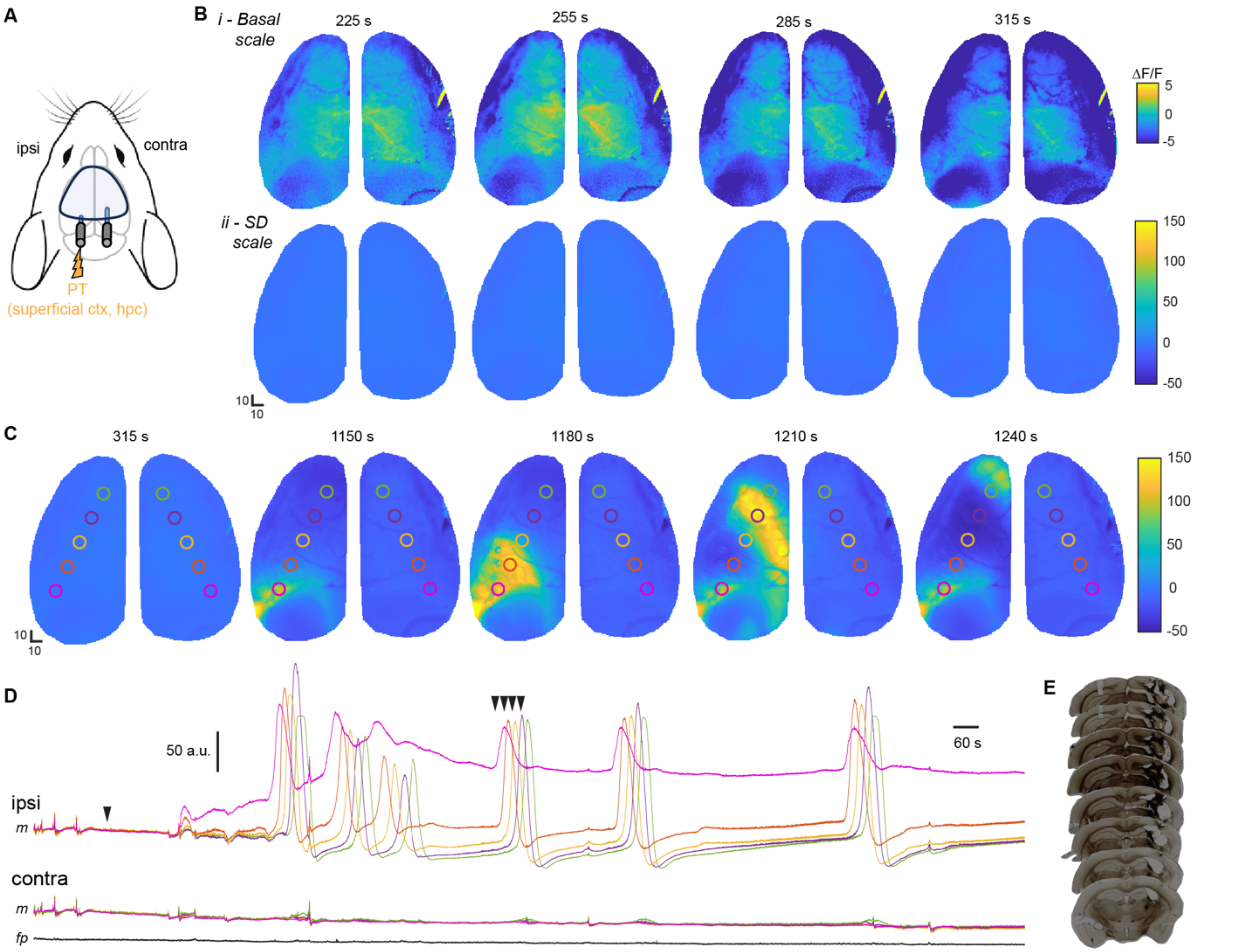


**Extended Data Figure 9. Unilateral hippocampal-neocortical photothrombosis triggers SDs restricted to ipsilesional neocortex. (A)** Schematic of mesoscale imaging window above superficial neocortex and bilateral fiberoptic implants (left: neocortex and dorsal hippocampus; right: dorsal hippocampus). **(B)** Baseline mesoscale neocortical GCaMP6f dynamics captured at ***(i)*** basal fluctuation and ***(ii)*** SD scales over 4 time points at 30 s intervals (see ΔF/F, right). **(C)** Representative mesoscale neocortical GCaMP6f images (basal: 315 s, SD: 1150-1240, 30 s intervals) highlighting an ipsilesionally-confined SD from ischemic neocortex. Photothrombosis (561 nm: 6 mW, 8 min) occurred at 600 s. **(D)** Representative structure/ROI-based (11 px$^2$, 35 µm/px) mesoscale imaging traces (n = 1, *m*) from female mouse for ROIs separated by 1.2 mm outward from the ischemic core in the SD propagation axis in ipsilesional (light, bold) neocortex, reflected in contralesional (dark) hemisphere following photothrombosis encroaching into neocortical tissue. Proximal ROI highlights TD, while distal ROIs indicate up to 5 cortical SDs confined to the ipsilesional neocortex and absent from the time-locked contralesional hippocampal photometry recording. **(E)** Coronal sections of representative female mouse brain experiencing focal neocortical/hippocampal photothrombosis, extracted 7 d post-illumination, indicates a unilateral stroke lesion encroaching on neocortex.

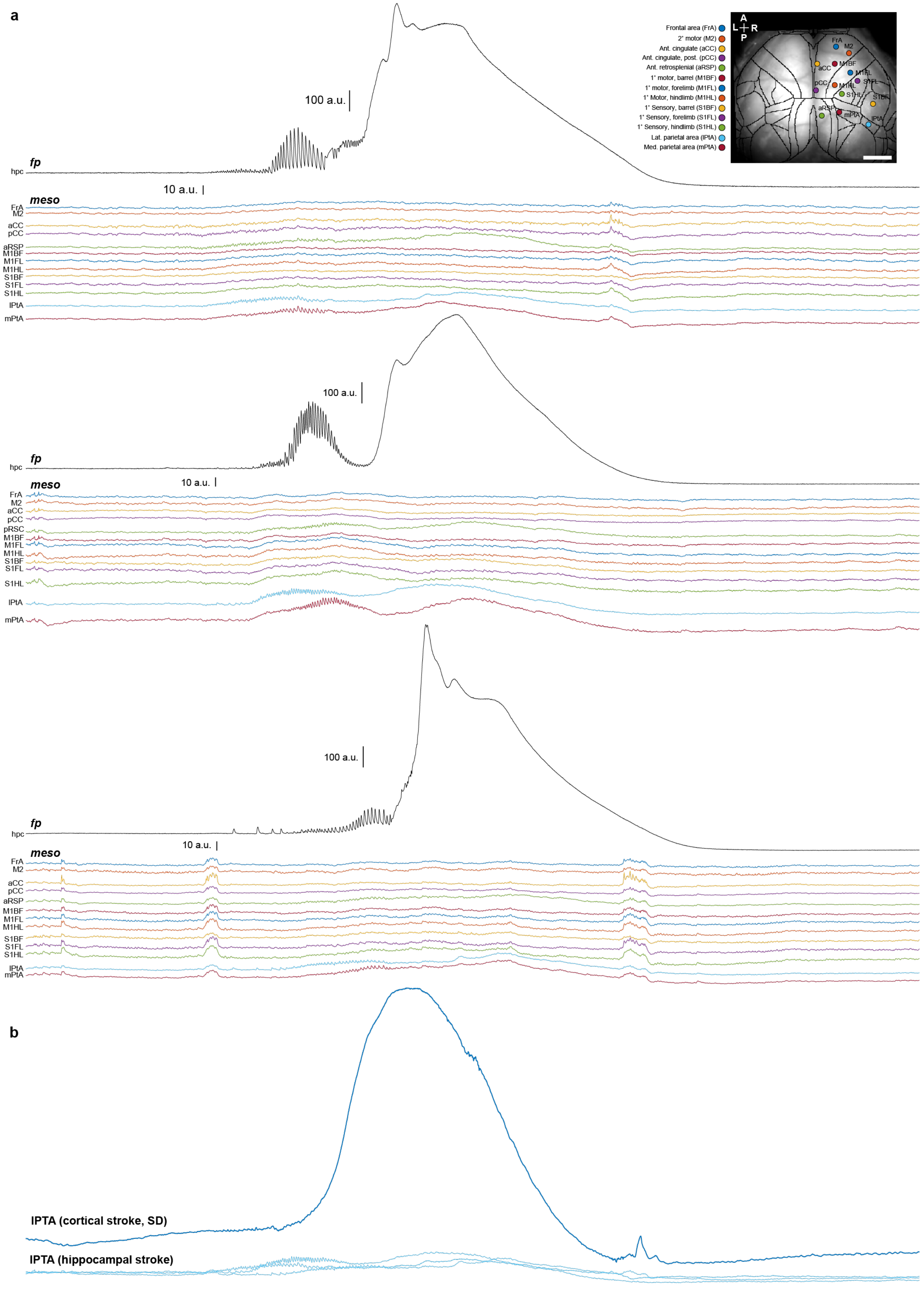
a
100 a.u.
fp
hpc
10 a.u.
meso
FrA
M2
aCC
pCC
aRSP
pRSC
M1BF
M1FL
M1HL
S1BF
S1FL
S1HL
lPtA
mPtA
Frontal area (FrA)
2° motor (M2)
Ant. cingulate (aCC)
Ant. cingulate, post. (pCC)
Ant. retrosplenial (aRSP)
1° motor, barrel (M1BF)
1° motor, forelimb (M1FL)
1° Motor, hindlimb (M1HL)
1° Sensory, barrel (S1BF)
1° Sensory, forelimb (S1FL)
1° Sensory, hindlimb (S1HL)
Lat. parietal area (lPtA)
Med. parietal area (mPtA)
A
L
R
P
b
lPTA (cortical stroke, SD)
lPTA (hippocampal stroke)

**Extended Data Figure 10. Cortical neurons demonstrate seizure-like $Ca^{2+}$ dynamics without SD during ischemia-induced contralesional hippocampal SD. (A)** 200 s epochs of contralesional recordings surrounding hippocampal SD (ΔF/F, fiber photometry) following unilateral hippocampal photothrombosis highlight the presence of rhythmic GCaMP6f dynamics in neocortical mesoscale imaging (ΔF/F) but the absence of a large slow component (cortical spreading depolarization) during hippocampal SD. **(B)** Contralesional mesoscale GCaMP6f traces from the lateral parietal area following hippocampal photothrombosis, normalized to the GCaMP6f response in the same region during a cortical stroke-induced SD.

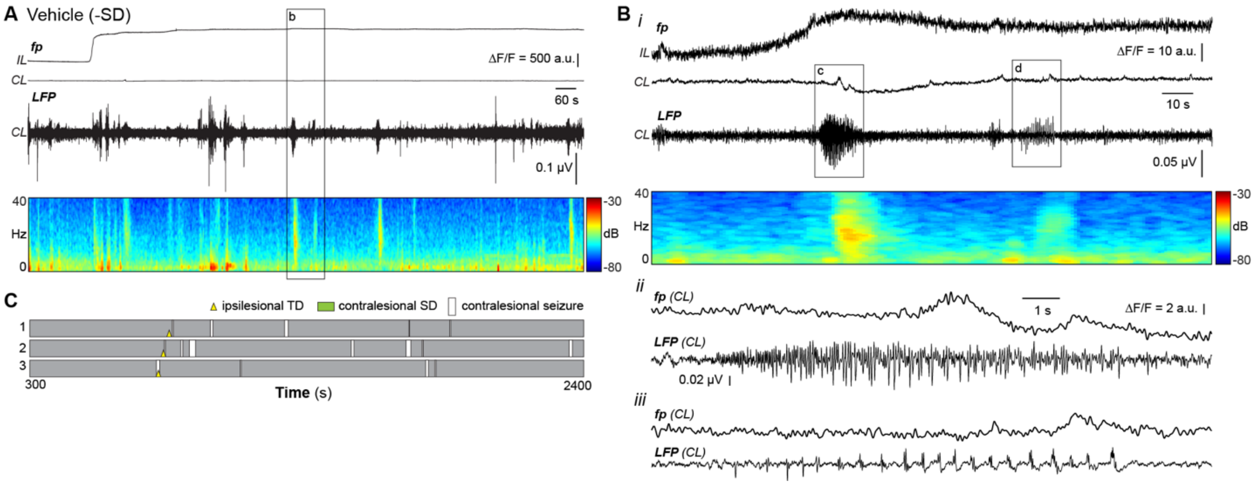


**Extended Data Figure 11. Contralesional electrographic seizures can occur in the absence of SD. (A)** Example ipsilesional (IL) and contralesional (CL) GCaMP6f (ΔF/F, red) paired with CL LFP (black trace) and time-varying frequency-power-time spectrogram in an awake, behaving female mouse (600 – 2400 s) pretreated with vehicle (PBS) via injection i.p. prior to baseline recording. **(B)** Window (180 s, *i*) surrounding representative seizures in the absence of SD. (***ii, iii***) Expanded sections (*15 s)* highlight differences in seizure waveform (LFP) relative to GCaMP6f dynamics. **(C)** Schematic overlay of contralesional recordings (LFP/FP) in vehicle-treated mice, highlighting incidence of electrographic seizure (white box) in relation to IL TD (yellow triangle), but absent of CL SD (green box).
